# Mapping RUNX2 transcriptional dynamics during multi-lineage differentiation of human mesenchymal stem cells

**DOI:** 10.1101/2023.03.30.534618

**Authors:** Kannan Govindaraj, Sakshi Kannan, Marcel Karperien, Janine N. Post

**Author notes:** KG and SK designed and performed the experiments and analyzed the data. JP conceived the idea and designed the experiments. MK critically revised the manuscript. All authors contributed to the writing of the manuscript. Authors declare no competing interests.

## Abstract

The multi-lineage differentiation capacity of human mesenchymal stem cells (hMSCs) enables its potential for tissue engineering and regenerative medicine. Master transcription factors play a key role during development, differentiation, homeostasis and disease pathology. RUNX2 is the master transcription factor for bone development, and it regulates several important signaling pathways during chondrogenic and osteogenic differentiation of hMSCs. However, modulation of RUNX2 activity during hMSC differentiation into various lineages is not yet fully described. We differentiated hMSCs into chondro-, osteo-, and adipogenic lineages and studied RUNX2 protein dynamics using Transcription Factor - Fluorescence Recovery After Photobleaching (TF-FRAP) at different time points. The TF-FRAP method can capture the dynamic changes of RUNX2 protein mobility at the single cell level resolution, and cluster analysis shows how RUNX2 dynamics change at subpopulation level in proliferating and differentiating hMSCs. Our data show that although whole hMSC population is exposed to differentiation stimuli, some subpopulations in hMSCs do not respond to environmental cues.

## Introduction

Human mesenchymal stem cells (hMSCs) are plastic, capable of self-renewal and multi-lineage differentiation. hMSCs are of great interest in tissue engineering due to its differentiation potential towards chondrogenic, osteogenic and adipogenic lineages. Treating osteochondral defects with allogenic or patients’ own MSCs have shown promising results (1). Several strategies, including MSC injection in the joint and implanting in vitro differentiated MSCs with or without supporting biomaterials, have been investigated (2). However, clinical translation of MSC-based therapies to treat osteochondral defects are not successful due to the functional mismatch between implanted hMSCs and chondrocytes or osteoblasts (3).

In vitro differentiation of hMSCs into chondrocytes results in either fibro cartilage or hypertrophic differentiation, and is thus not suitable for the treatment of cartilage defects. Osteogenic differentiated hMSCs lack the mechanical properties of osteoblasts, thereby limiting successful bone implants (4). Despite much progress, complete knowledge of the signaling interplay during MSC differentiation into multiple lineages is still lacking. Exploring the intra- and extracellular signaling network involved in MSC differentiation will aid in advancing tissue-engineering strategies and expand our knowledge in MSC lineage commitment (5).

The role of transcription factors is paramount in lineage commitment and differentiation. RUNX2 (6), SOX9 (7) and PPAR*γ* (8) are the master transcription factors for osteoblast, chondrocyte, and adipocyte differentiation, respectively. RUNX2, the master regulator of bone development, plays a key role in multiple signaling pathways associated with bone development, including TGF, BMP and WNT (9, 10). RUNX2 directly regulates the expression of extracellular protein, COL10A1, which is abundantly present in the bone (11). In addition, RUNX2 plays an essential role during chondrogenic differentiation of hMSCs and hypertrophic differentiation of chondrocytes (12). The role of RUNX2 is minimal and its activity is inhibited during adipogenic differentiation of hMSCs (8). Although much is described about RUNX2 activity during various lineage differentiation of hMSCs, its activity has been studied by quantifying target gene or protein expression levels, using static methods such as qPCR and western blot, respectively. However, hMSCs are heterogenic and contain several subpopulations of cells (13–15). Studying the RUNX2 protein activity at the single cell level will help us to understand how its transcriptional activity changes during differentiation at the subpopulation level.

We have previously shown how SOX9 protein dynamics change among the subpopulation of hMSCs during chondro-, osteo- and adipogenic differentiation (16). In this report, using TF-FRAP, we have studied how RUNX2 protein dynamics changes during chondro-, osteo- and adipogenic differentiation of hMSCs.

## Materials and Methods

### Cell Culture and Transfection

hMSCs were isolated from bone marrow from patients with no known musculoskeletal diseases. hMSCs were cultured in MEM (22571-038, Gibco), supplemented with 10% FBS (F7524, Sigma), 200mM glutaMax (35050-38, gibco), 20mM ascorbic acid 2 phosphate (AsAP, Sigma, A8960), 100 ng/ml bFGF (Neuromics, RP80001), and 100 U/ml of Penicillin-Streptomycin (15140122, gibco) at 37°C with 5% CO_2_. hMSCs were expanded and used within 4 passages. C20/A4 cells were cultured in DMEM (Gibco, USA) with 10% FBS. Human primary chondrocytes were cultured in chondrocyte proliferation media containing DMEM supplemented with 10% FBS, 20mM ascorbic acid 2 phosphate (AsAP, Sigma, A8960), L-Proline (40 *µ*g/ml) and non-essential amino acids, at 37°C with 5% CO_2_. For transfection, cells were seeded on a sterile glass coverslip (40,000 per well) placed inside the well. eGFP-RUNX2 plasmid was kindly given by Gary S Stein (17). Lipofectamine 3000 with P3000 Reagent (Life Technologies, USA) was used for transfection of hMSCs and Lipofectamine LTX with Plus reagent was used for transfection of C20/A4 cells and hPCs and the manufacturer’s protocol was followed.

### Cell cycle synchronization

hMSCs were seeded on microscopic coverslip placed inside a 24-well plate in hMSC proliferation media. Next day, cells were transfected as described above and maintained in proliferation media with serum for about 8 hours to allow cells to produce eGFP-RUNX2 protein. For TF-FRAP: Transfected cells were maintained in serum free proliferation media 24 hours prior to TF-FRAP measurements. During TF-FRAP measurements, cells were maintained in the imaging buffer as described below. For imaging eGFP-RUNX2 nuclear localization pattern: for 24+0h time point, transfected cells were maintained in serum free proliferation media 24 hours prior to imaging. For 24+6h time point, post 24h of serum starvation, cell cycle was again started by replacing the media with proliferation media with serum. Cells were imaged 6h after the start of cell cycle, so the time point was 24+6h. Cells in two different coverslips were used for 24+0h and 24h+6h time points study. During imaging, cells were maintained in the imaging buffer, at 37°C.

### hMSC differentiation

hMSCs were differentiated into chondrogenic, osteogenic and adipogenic lineages by culturing them in the respective differentiation media from day 0. Culture media were refreshed every 3 – 4 days. Chondrogenic differentiation medium: hMSCs (5,000 cells/cm^2^) were cultured in DMEM (Gibco) supplemented with 1% of Insulin-Transferrin-Selenium (ITS) mix (41400045, Gibco), 40 *µ*g/ml of L-proline, 50 *µ*g/ml of AsAP, 1% of sodium pyruvate (S8636, Sigma), 100 U/ml of Penicillin-Streptomycin, and freshly added TGF1 (10 ng/ml, 7754-BH, RD Systems) and 10^*−*7^ M dexamethasone (Dex, D8893, Sigma). Osteogenic differentiation medium: hMSCs (1,000 cells/cm^2^) were cultured in MEM supplemented with 10% FBS, 100 U/ml of Penicillin-Streptomycin, 200mM GlutaMax, 50 *µ*g/ml AsAP and freshly added Dex (10^*−*8^ M). Adipogenic differentiation medium: hMSCs (10,000 cells/cm^2^) were cultured in MEM supplemented with 10% FBS, 100 U/ml of Penicillin-Streptomycin, 200mM glutaMax, 50 *µ*g/ml AsAP, freshly added Dex (10^*−*6^ M), 10 *µ*g/ml Insulin (I9278, Sigma), 0.5 mM IBMX (I5879, Sigma) and 0.2 mM Indomethacin (57413, Sigma).

### Imaging Buffer

Imaging was performed in Tyrode’s buffer (18) with freshly added 20 mM glucose (Gibco) and 0.1% BSA (Sigma). Tyrode’s buffer is composed of 135 mM NaCl (Sigma), 10 mM KCl (Sigma), 0.4 mM MgCl_2_ (Sigma), 1 mM CaCl_2_ (Sigma), 10 mM HEPES (Acros organics), pH adjusted to 7.2, filter sterilized (0.2 *µ*m) and stored at -20°C.

### Transcription Factor - Fluorescence Recovery After Photo-bleaching

Proliferating or differentiating hMSCs were plated on poly-l-lysine (0.01%, Sigma) coated glass cover slips 2 days before transfection and were transiently transfected with eGFP-RUNX2 a day before FRAP experiments. Chondrogenically differentiating hMSCs did not attach on the coverslip without serum in the differentiation media. So, only for chondrogenic differentiation, serum was added to differentiation media, when we reseeded cells on coverslips for TF-FRAP, again media was replaced without serum, once the cells were attached. FRAP was performed at four time points, i.e., 0 day (proliferation), day 2, day 8 and day 15 or 23, for all the three lineages. Cells were maintained in imaging buffer during FRAP measurements and the mobility of transcription factors was measured at the steady-state (i.e. without other stimuli or FBS added). FRAP measurements were performed using a Nikon A1 laser scanning confocal microscope (Nikon, Japan). with 60X/1.2 NA water immersion objective, 488 nm Argon laser at 0.50% (0.12 *µ*W at the objective) laser power for eGFP-RUNX2. The temperature was maintained at 37°C with an OkaLab temperature controller. A frame size of 256×256 pixels covering the whole nucleus was scanned. The pixel size was 0.12 *µ*m. A representative circular region of 2.9 *µ*m diameter was bleached with one iteration (60 ms) of 50% (34.3 *µ*W) laser power. Twenty-five pre-bleach images were taken and the last 10 pre-bleach fluorescence intensity values were averaged to normalize the post-bleach fluorescence recovery curve. After bleaching, imaging was performed at 4 frames/sec for 60 sec post-bleach. FRAP experiments were performed on at least 40 cells per condition. To assess the statistical significance between the conditions Mann-Whitney U tests were applied using Origin software. Matlab™ was used to analyze the FRAP data and the script is available upon request. A diffusion uncoupled, two-component method was used to interpret our FRAP results as mentioned in (19).

### Clustering and statistical analysis

We applied unsupervised hierarchical clustering to cluster RUNX2 dynamics data. TF-FRAP variables, such as Immobile Fraction, Recovery half-time of A_1_ and A_2_ were used as input for cluster analysis. The following clustering parameters were used: Distance type: Euclidean, Cluster method: Furthest neighbor, and Find Clustroid: Sum of distances. The distinct shapes of FRAP curves served as good references to identify and determine the number of clusters in our FRAP data. In some cases, two clusters as segregated by unsupervised hierarchical clustering, had very close similarity with negligible differences (statistically insignificant) in the FRAP rates. Therefore, those clusters were combined and considered one cluster. Clusters within a differentiation lineage at different time points that showed no significant difference (at P*≥*0.05, Mann-Whitney U-test) for at least two FRAP variables, were considered as same clusters, and were given the same cluster name/identifier. OriginPro (Origin Labs, USA) was used for clustering analysis.

### Generation of heat-maps

For convenience, clusters as segregated by hierarchical clustering were reordered per time-point within the differentiation lineage in the ascending order of immobile fraction values. All three FRAP variables, IF, recovery half-time (t1/2) of A_1_ and t1/2 of A_2_ were normalized between 0 and 100. Within a variable, the lowest and highest values were set to 0 and 100 respectively. OriginPro was used to make heat-maps.

### Immunofluorescence

hMSCs cultured on microscopy coverslips were fixed for 10 min with freshly prepared 4% Paraformaldehyde in PBS, pH7.2. Cells were washed with ice cold PBS, 3 times with 5 min interval. Cells were blocked and permeabilized for 15 mins with blocking solution containing 1 mg/ml BSA and 0.1% Triton X-100 in PBS. Cells were incubated with mouse anti-human RUNX2 primary antibody (ab76956, Abcam) in the blocking solution for 30 min at 37°C. Cells were washed with PBS, 3 times with 5 min intervals. Cells were incubated with goat anti-mouse AF-647 antibody (A-21235, Invitrogen) in the blocking solution (with 0.05% Triton X-100) for 30 min at 37°C. Cells were washed with PBS, 3 times with 5 min interval. Cell were stained with DAPI (1:100 dilution to 5 ng/ml) for 5 min in PBS. Cells were washed with PBS, 3 times with 5 min interval and mounted to microscopy glass side using Vecta shield mounting medium (Vector Laboratories). Cells were imaged using a Nikon A1 Confocal microscope with 60x water immersion objective with 1.2 NA.

### Histological Staining

Differentiation of hMSCs into chondrogenic, osteogenic and adipogenic lineage is followed by Alcian blue, ALP and Oil Red O staining, respectively, at various time points. Alcian blue staining was used to stain GAG produced by chondrogenic differentiated hMSCs at day 0, day 2, day 8 and day 23. Cells were fixed with 10% buffered formalin (HT501128, Sigma) for 15 mins and washed twice with ice cold PBS. Cells were incubated with freshly prepared Alcian Blue 8GX (A3157, Sigma) staining solution (0.5% w/v in 1M HCl, pH 1.0) for 30 mins and the cells were rinsed with dH2O until the staining solution is washed off. ALP staining was used to stain alkaline phosphatase activity by osteogenic differentiated hMSCs at day 0, day 2, day 8 and day 14. ALP staining kit (S5L2-1KT, Sigma) was used for the staining and the manufacturer’s protocol was followed. Oil Red O staining was used to stain neutral triglycerides and lipids produced by adipogenic differentiated hMSCs at day 0, day 2, day 8 and day 23. Cells were fixed with 10% buffered formalin for 15 mins and washed twice with ice cold PBS and dH20. Cells were incubated in 60% isopropanol for 5 mins and then in freshly filtered Oil Red O staining solution for 5 mins and the cells were rinsed with dH2O until the staining solution was washed off. Cells were observed using a Nikon ECLIPSE TE300 microscope with 4x (0.13 NA) objective for all the three staining.

## Results

### Interpretation of TF-FRAP data

TF-FRAP data is interpreted by applying an appropriate model, representing actual events contributing to fluorescence recovery. An optimal TF-FRAP model is based on the number of events and time-scale in which the event occurs. In case of transcription factors, three types of events are possible, i.e., unbound transcription factors freely moving (diffusing) inside the nucleus, transcription factors weakly bound to DNA (fast moving) and strongly bound/exchanging at its binding site on DNA (slow moving) (20). Diffusion is an extremely fast event, occurs even before capturing the first post-bleach image and thus, is not included in the model. Therefore, a diffusion uncoupled, two-component fit method is one of the models suitable for interpreting the TF-FRAP data.

In this model, the first component indicates the fast moving (A_1_) fraction, the second component indicates the slow moving (A_2_) fraction of transcription factors, and these two events occur in distinct time-scales (21). Dynamic rates of a transcription factor, such as recovery half-times, diffusion constants of A_1_ and A_2_ and Immobile Fraction (IF) can be calculated from the TF-FRAP curve. For detailed information, we refer the readers to (19–21) We have previously shown that SOX9 binding to DNA determines its transcriptional activity (19). Higher immobile fraction (DNA binding) of SOX9 and or longer recovery half-time of A_2_ increased its target gene expression and vice versa.

### Cellular heterogeneity leads to diverse RUNX2 mobility patterns

RUNX2 transcriptional activity and its target genes expression levels are known to be dynamically regulated during various lineage differentiation (22). Since the activity of a transcription factor is coupled to its dynamics (19), to understand RUNX2 activity, we measured changes in RUNX2 mobility during chondrogenic, osteogenic and adipogenic differentiation of hMSCs. RUNX2 activity is known to change dynamically during hMSC differentiation (reviewed in (11)). For example, RUNX2 activity increased during osteogenic (10) and initial stages of chondrogenic differentiation (23), while it decreased during adipogenic differentiation (24). Although these studies were based on RUNX2 or its target gene or protein expression levels, we expected changes in RUNX2 protein dynamics in line with these reports. However, data analysis by averaging FRAP values per time point did not yield any useful information, except that the RUNX2 mobility increased during hMSC differentiation into any lineage (Figure S1). Interestingly, we observed diverse mobility patterns of RUNX2 in proliferating and differentiating hMSCs.

To gain more insight into RUNX2 transcriptional activity at the sub population level, we clustered the FRAP data using an unsupervised hierarchical clustering method. Cluster analysis revealed the presence of at least four clusters with distinct dynamic rates in RUNX2 FRAP data in undifferentiated hMSCs (Figure S2 and Table S1). To check whether these diverse mobility patterns are inherent property of any cell type, we measured RUNX2 mobility in the C20/A4 cell line and in human primary chondrocytes. Diversity of RUNX2 mobility was lower in the C20/A4 cells, whereas we observed at least two clusters with distinct mobility patterns in hPCs (Figure S3).

To rule out the possibility of cell cycle playing a role in generating the diverse mobility patterns, we measured RUNX2 mobility in 24-h cell cycle synchronized hMSCs. We observed four clusters in the synchronized hMSCs as well (Figure S4). Although hMSC doubling time is known to be 30-39h (25), we could synchronize transfected cells only for 24 h, as they start to die afterwards. In addition, we also observed diverse RUNX2 mobility in chondrogenically differentiating hMSCs, where cell division is limited due to lack of serum. Moreover, RUNX2 dynamics data from other cell types (C20/A4 and hPCs) and heterogenic nature of hMSCs indicate that the diverse mobility patterns of RUNX2 in hMSCs are due to cellular heterogeneity of hMSCs. Thus, we applied unsupervised hierarchical clustering to cluster the cells in further data analysis. These data are consistent with our SOX9 dynamics study, which also showed at least four clusters in undifferentiated hMSCs with distinct FRAP rates.

### RUNX2 binding and its residence time on DNA decrease during chondrogenic differentiation

RUNX2, along with SOX9, is known to co-ordinate chondrogenic differentiation of hMSCs (26, 27). To further understand the role of RUNX2 protein during chondrogenic differentiation, we measured RUNX2 dynamics by TF-FRAP in chondrogenically differentiating hMSCs at day 2, 8 and 15. Since RUNX2 plays a role in the initial stages of chondrogenic differentiation (23), we expected higher immobile fraction and/or longer recovery half-times in the early time-points of chondrogenically differentiating subpopulations.

Unsupervised hierarchical clustering showed at least six clusters with distinct FRAP rates at day 2, and five clusters at day 8 and 15. If two out of three RUNX2 dynamic rates of a cluster was not significantly different among any of the clusters of chondrogenically differentiating hMSCs (including undifferentiated hMSCs), they were considered same cluster and given same cluster name. Heat maps of FRAP data show the changes in RUNX2 dynamics at the single cell level during chondrogenic differentiation (Figure 1A-D). Several new clusters with distinct RUNX2 dynamics appeared during chondrogenic differentiation. Clusters J, K and L were present at day 8 and 15. Day 2 showed five new clusters, indicating RUNX2 interaction with DNA changes considerably during in the initial phase of chondrogenic differentiation. TF-FRAP curves of RUNX2 during chondrogenic differentiation at individual cluster level, and relationship between the IF and half-time to recover of A_2_ are shown in figure S5. Percentage of cells per cluster is shown in figure S8A and B. GAG staining confirmed chondrogenic differentiation of hMSCs (Figure 1E).

**Fig. 1.**
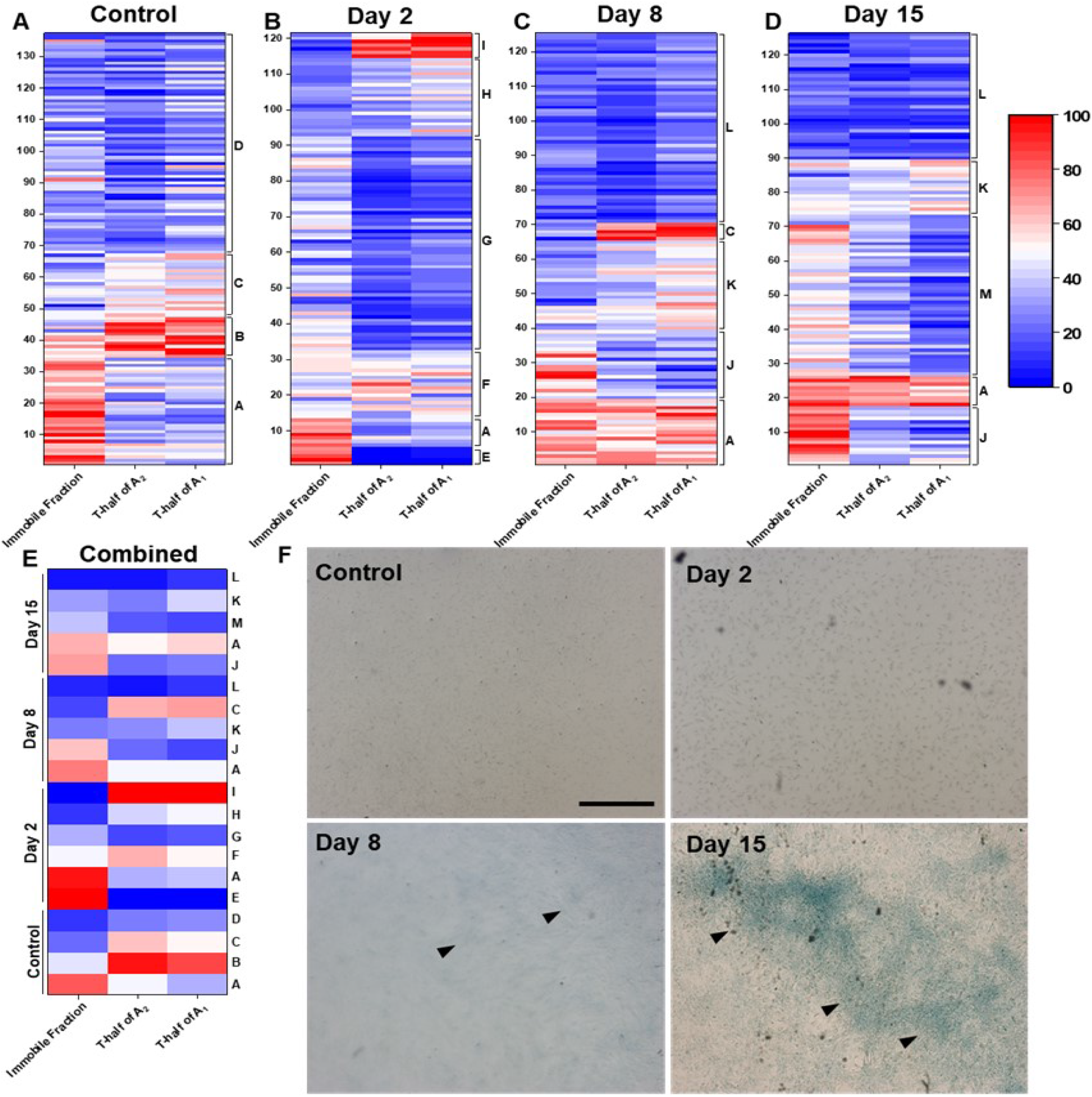
RUNX2 binding and residence time on DNA decrease during chondrogenic differentiation of hMSCs. RUNX2 mobility was measured at day 0 (undifferentiated), 2, 8, and 15 of chondrogenic differentiation. Unsupervised hierarchical clustering showed four clusters in undifferentiated hMSCs (A), six clusters in day 2 (B) and five clusters in day 8 and 15 (C and D). (E) Heat map showing averaged FRAP values per cluster across time-points, letters at the right indicate cluster ID. Although several new clusters with distinct RUNX2 dynamics appeared during chondrogenic differentiation, cluster A of undifferentiated hMSCs appeared in all three time points of chondrogenically differentiating hMSCs. Three hMSC donors were used for this study. Combined data from all donors is shown here. Three FRAP variables, IF, T-half of A2 and A1, were used for cluster analysis. FRAP variables were normalized (0, 100) to make heat maps. N*≥*121, per time point. (F) Increased Alcian blue staining at later time points confirms the chondrogenic differentiation of hMSCs. Black arrow heads indicate GAG staining. Scale bar: 500*µ*m.

At day 2, immobile fraction increased in two clusters (E and A, > 60%) and recovery half-times in four clusters A, F, H and I (>1.6 sec for A_1_ and >10 sec for A_2_) as comparted to day 8 and 15 (Figure 1E, S5and Table S2). Except clusters A and C (at day 8 and/or 15), recovery half-time of A_2_ was lower than 10 sec as compared to higher recovery half-time of A_2_ in clusters at day 2. A new cluster named L, which was present at day 8 and 15, had lower IF (25%) and shorter recovery half-times of A_1_ and A_2_ (0.8 sec and 4.5 sec, respectively). Interestingly, cluster G, which contained 50% of cells at day 2, had comparatively lower recovery half-time of A_2_, next to cluster E. Cluster L, which contained 44% of cells at day 8 and 29% of cells at day 15, had lowest IF and recovery half-times at those time points. Clusters A and J at day 8 and 15, which had higher IF and/or longer recovery half-times as compared to other clusters within their time-point, comparatively lower number of cells (16% or less). This suggests that RUNX2 activity decreased during the later stages (day 8 and 15) of chondrogenic differentiation as compared to day 2 and undifferentiated hMSCs (Figure 1E, and Table S2). Recovery half-times of clusters in chondrogenically differentiating clusters were higher than adipogenic differentiation, but lower than osteogenic differentiation (Figure S15).

Although heat maps show dynamics data at the single cell level, this change is not clearly visible as the normalization of FRAP data is influenced by the highest and lowest values in the data set, making it hard to compare rates between different time points. During differentiation, there could be different scenarios for the efficiency of cell differentiation. For example, one can assume that the transcription factor RUNX2 responds similarly to the differentiation stimuli in all cells, in which case we would expect fewer clusters in the later time-points. In another scenario, cells in one cluster may not respond to the differentiation stimuli, while cells in other clusters differentiate at different rates. In this case, one could expect (some of) the clusters from the undifferentiated cells, and multiple new clusters as differentiation progresses. That is, if the activity of the transcription factor is indeed dependent on the stage of differentiation. By comparing the rates of the different clusters over the different differentiation time-points, we investigated what clusters were similar to the clusters found in undifferentiated cells. Interestingly, cluster A, as identified in undifferentiated hMSCs, was present at all three time points, albeit in low numbers. This indicates that at least one subpopulation of cells was not affected by the differentiation stimuli.

### RUNX2 residence time increases during osteogenic differentiation

RUNX2 is one of the driving factors of osteogenic differentiation (10). To understand its transcriptional dynamics during osteogenic differentiation, we measured its dynamics by TF-FRAP at day 2, 8 and 23 in osteogenically differentiating hMSCs. Heat maps show RUNX2 dynamics at the single cell level in control (undifferentiated) and at day 2, 8 and 23 (Figure 2A-D). During osteogenic differentiation, we observed six new clusters (E – J) and cluster B from undifferentiated hMSCs was present in all three time points, indicating that this cluster did not respond to the differentiation stimuli. Cluster C was also present at day 2, suggesting cells in this cluster are slow to respond to the differentiation stimuli.

**Fig. 2.**
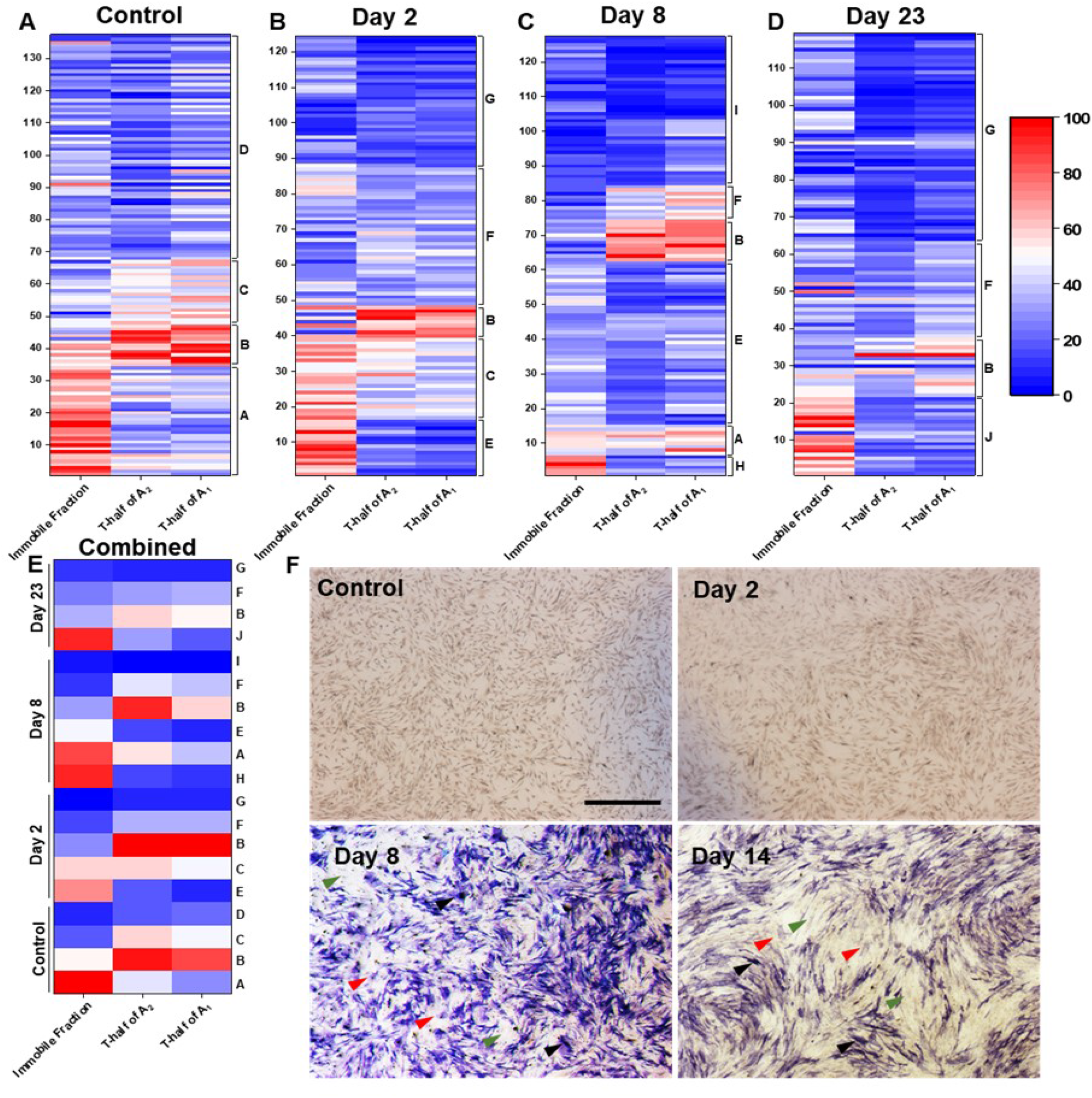
RUNX2 binding and residence time on DNA increase during osteogenic differentiation of hMSCs. RUNX2 mobility was measured at day 0 (undifferentiated), 2, 8, and 23 of osteogenic differentiation. Unsupervised hierarchical clustering showed five, six and four clusters at day 2, 8 and 23 of osteogenically differentiated hMSCs, respectively (A-D). (E) Heat map showing averaged FRAP values per cluster across time-points, letters at the right indicate cluster ID. Cluster B from undifferentiated hMSCs and a new cluster F in differentiating hMSCs appeared at all three time points. Three TF-FRAP variables IF, T-half of A2 and A1 were used for cluster analysis. TF-FRAP variables were normalized (0, 100) before making heat maps. Three hMSC donors were used for this study. Combined data from all donors are shown here. N*≥*119, per time point. (F) Increased ALP staining in the later time points confirm the osteogenic differentiation of hMSCs. Black arrow heads point cells with very high ALP production in some cells, red arrow heads point cells with lower ALP production and green arrow heads points cells with now ALP production. This indicates that subpopulation cells in differentiating hMSCs respond differently to the differentiation stimuli. Scale bar: 500*µ*m.

Since RUNX2 is a bone specific transcription factor, we expect it to be more active during osteogenic differentiation as compared to chondrogenic differentiation. Higher transcriptional activity results in a higher immobile fraction and longer t-half times [(9) As expected, of the six new clusters in the osteogenic differentiation, two clusters (F and J) showed longer recovery half-time of A_2_ (10 sec, Table S4) as compared to the those in the other two lineages (Figure S15, Table S3 and S5). A new cluster, F, was present at all three time-points, with recovery half-times of 10.35 sec (A_2_) and 1.67 (A_1_). IF and recovery half-times were higher during osteogenic differentiation as compared to undifferentiated hMSCs (Figure 2E). In addition, clusters C and B from undifferentiated hMSCs with longer recovery half-times were also present during osteogenic differentiation. However, contrary to our expectation, the IF was lower in clusters of osteogenic differentiation as compared to other two differentiation lineages (Figure S15 and Table S3-S5).

TF-FRAP curves and relationship between IF and half-time to recover of A_2_ are presented in figure S6A - F. The number of cells per cluster are presented in figure S8. ALP staining confirms osteogenic differentiation of hMSCs (figure 2E). Some cells produced very high amounts of ALP (indicated by black arrowhead), some cells produced comparatively low levels of ALP (indicated by red arrowhead), while others did not visibly produce ALP (indicated by green arrow head). This suggests that, although all cells were exposed to same differentiation stimuli, sub populations of cells in hMSCs respond to these stimuli with varying degrees, some produce ALP at high levels and some at low levels, while others do not visibly respond. Assuming that the levels of ALP production are correlated to the osteogenic differentiation, these varying levels of differentiation indicate an increase in RUNX2 transcriptional activity and thus in a decrease in RUNX2 dynamics, and vice versa.

### RUNX2 residence time decrease during adipogenic differentiation

We measured RUNX2 dynamics by TF-FRAP in adipogenically-differentiating hMSCs to understand its transcriptional activity. TF-FRAP rates at the single cell level during adipogenic differentiation at day 0 (undifferentiated), 2, 8 and 23 are shown in the heat maps (Figure 3A-D). Along with eight new clusters (E – L) with distinct RUNX2 dynamics, cluster C from undifferentiated hMSCs were present in the day 2 and 23 of adipogenically differentiating hMSCs. Although cluster C was not present at the day 8, cluster G at this time point closely resembled to cluster C. However, the FRAP rates were significantly different from cluster C, so we consider it as a separate new cluster. This suggests the cluster C did not respond to the adipogenic differentiation stimuli.

**Fig. 3.**
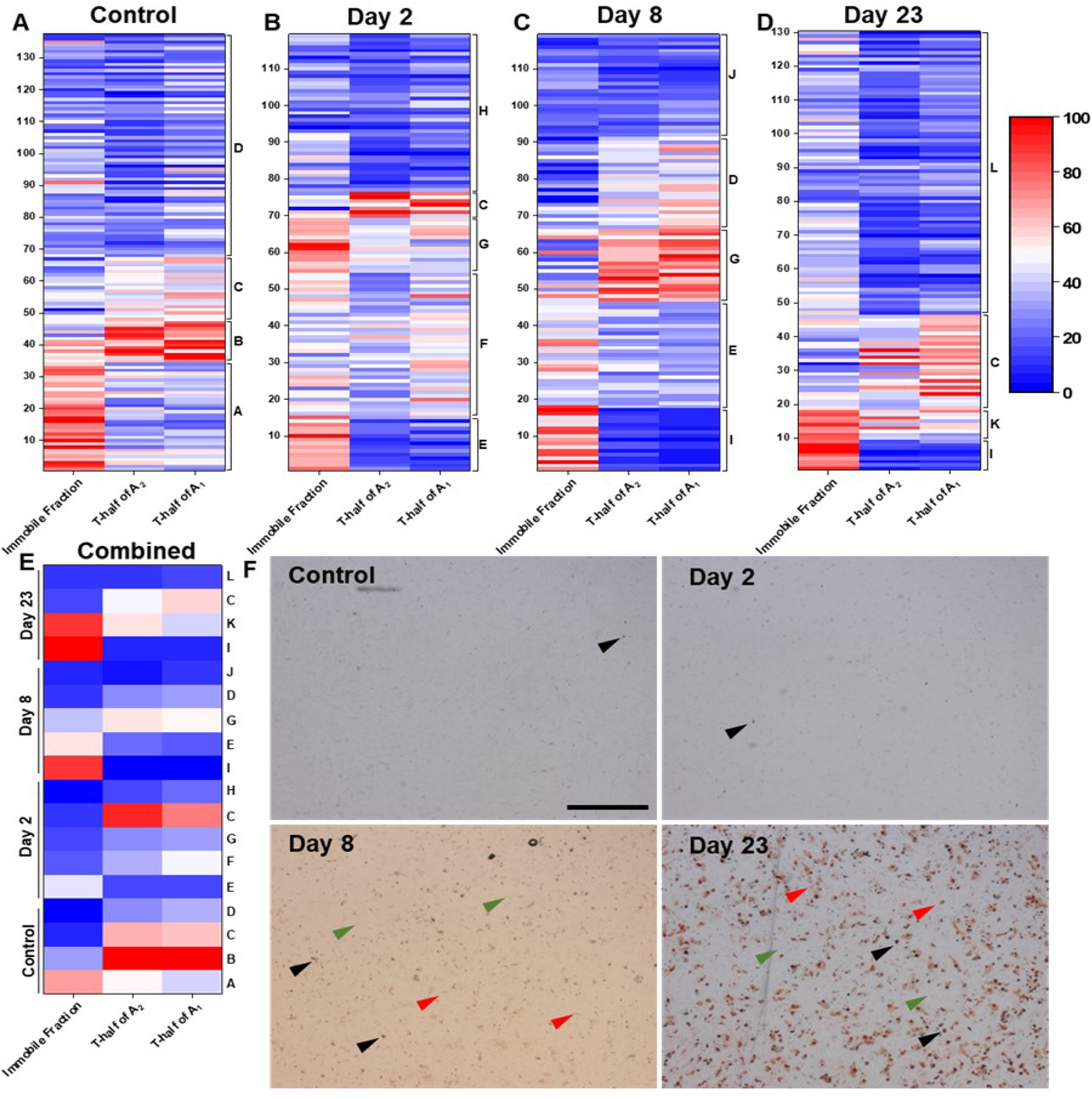
RUNX2 residence time on DNA decrease during adipogenic differentiation of hMSCs. eGFP-RUNX2 mobility was measured at day 0 (undifferentiated), 2, 8, and 23 of adipogenic differentiation. Unsupervised hierarchical clustering showed five clusters at day 2 and 8 and four clusters at day 23 in adipogenically-differentiated hMSCs. Along with several new clusters, clusters C or D from undifferentiated hMSCs also appeared during the adipogenic differentiation. (E) Heat map showing averaged FRAP values per cluster across time-points, letters at the right indicate cluster ID. Three FRAP variables IF, T-half of A2 and A1 were used for cluster analysis. FRAP variables were normalized (0, 100) before making heat maps. Three hMSC donors were used for this study. Combined data from all donors are shown here. N*≥*119, per time point. (F) Increased Oil Red O staining over time points confirm adipogenic differentiation of hMSCs. Black arrow heads indicate the cells with very high lipid production, red arrow heads indicate cells with lower lipid production and greed arrow heads indicate cells with no lipid production. This indicates subpopulation of cells in differentiating hMSCs do not respond similarly to the differentiation stimuli. Scale bar: 500*µ*m.

The role of RUNX2 is minimal in adipogenic differentiation of hMSCs (24), and we expected higher RUNX2 mobility (lower IF and shorter recovery half-times) in adipogenically differentiated cells. We found that most new clusters (E, F, G, H, I, J and L) were more highly mobile (recovery half-times of A_2_, <10 sec), while clusters G and K were less mobile with >10 sec of recovery half-time of A_2_. Moreover, recovery half-times of many clusters were comparatively shorter than chondrogenically (Table S2) and osteogenically (Table S3) differentiated hMSCs (Figure S15). TF-FRAP curves of individual clusters and their relationship between IF and recovery half time of A_2_ are shown in figure S7 and TF-FRAP values are presented in the table S4.

The percentage of cells per cluster are shown in figure S8D. Interestingly, clusters H, J and L (of day 2, 8 and 23, respectively), which contained comparatively higher percentage of cells within their time points, showed comparatively lower IF (33%) and shorter recovery half-times of A_2_ (5.68 sec) and A_1_ (0.96 sec, table S4). This suggests RUNX2-DNA binding (IF) and its residence time (recovery half-times) were drastically reduced during adipogenic differentiation (Figure 1E). Cluster I (of day 8 and 23) had higher immobile fractions (65%), but lower recovery half-times of A_2_ (4.38 sec) and A_1_ (0.56 sec), indicating little active exchange of RUNX2 with DNA (Table S4).

Oil Red O staining at day 8 and 23 show varied degrees of oil droplets among adipogenically differentiating cells (Figure 3E). Some cells contained very high amount of lipid droplets (indicated by black arrow heads) and some cells with low lipid droplets (indicated by red arrow heads) and some cells without any lipid droplets (indicated by green arrow heads). These data again suggest that, although all cells were exposed to same differentiation stimuli, sub population of cells respond differently resulting in distinct RUNX2 dynamics.

### RUNX2 dynamics of subpopulation of cells in hMSCs match with hPCs

hMSCs differentiate into articular chondrocytes and form cartilage tissue, which helps the smooth movement of joints (28). In case of OA, healthy cartilage tissue is damaged and chondrocytes differentiate into hypertrophic chondrocytes. During cartilage homeostasis, RUNX2 activity is minimal and its activity is increased during hypertrophic differentiation (29). To get more insights into this differentiation process, we compared RUNX2 dynamics of undifferentiated hMSCs with healthy hPCs and chondrogenically differentiated hMSCs (day 15) with healthy and OA hPCs.

hPCs themselves had two subpopulations of cells (we named them cluster 1 and 2) with distinct RUNX2 dynamics. We expected that RUNX2 dynamics of healthy hPCs might match, or be similar to, clusters in undifferentiated hMSCs that are expected to be chondro progenitors. Interestingly, the TF-FRAP rates of cluster 2 of healthy hPCs were similar to those of cluster C (changes were statistically insignificant in at least two TF-FRAP variables) of undifferentiated hMSCs, suggesting this cluster may be one of the progenitor subpopulations for chondrogenic differentiation.

We also compared RUNX2 dynamic rates of chondrogenically-differentiated hMSCs (day 15) with healthy and OA hPCs. We expected that if chondrogenically-differentiated hMSCs are identical to healthy or OA hPCs, their RUNX2 dynamics would be similar. Interestingly, cluster 1 of healthy and OA hPCs had similar RUNX2 FRAP rates to that of cluster J of chondrogenically-differentiated hMSCs (Figure 4).

**Fig. 4.**
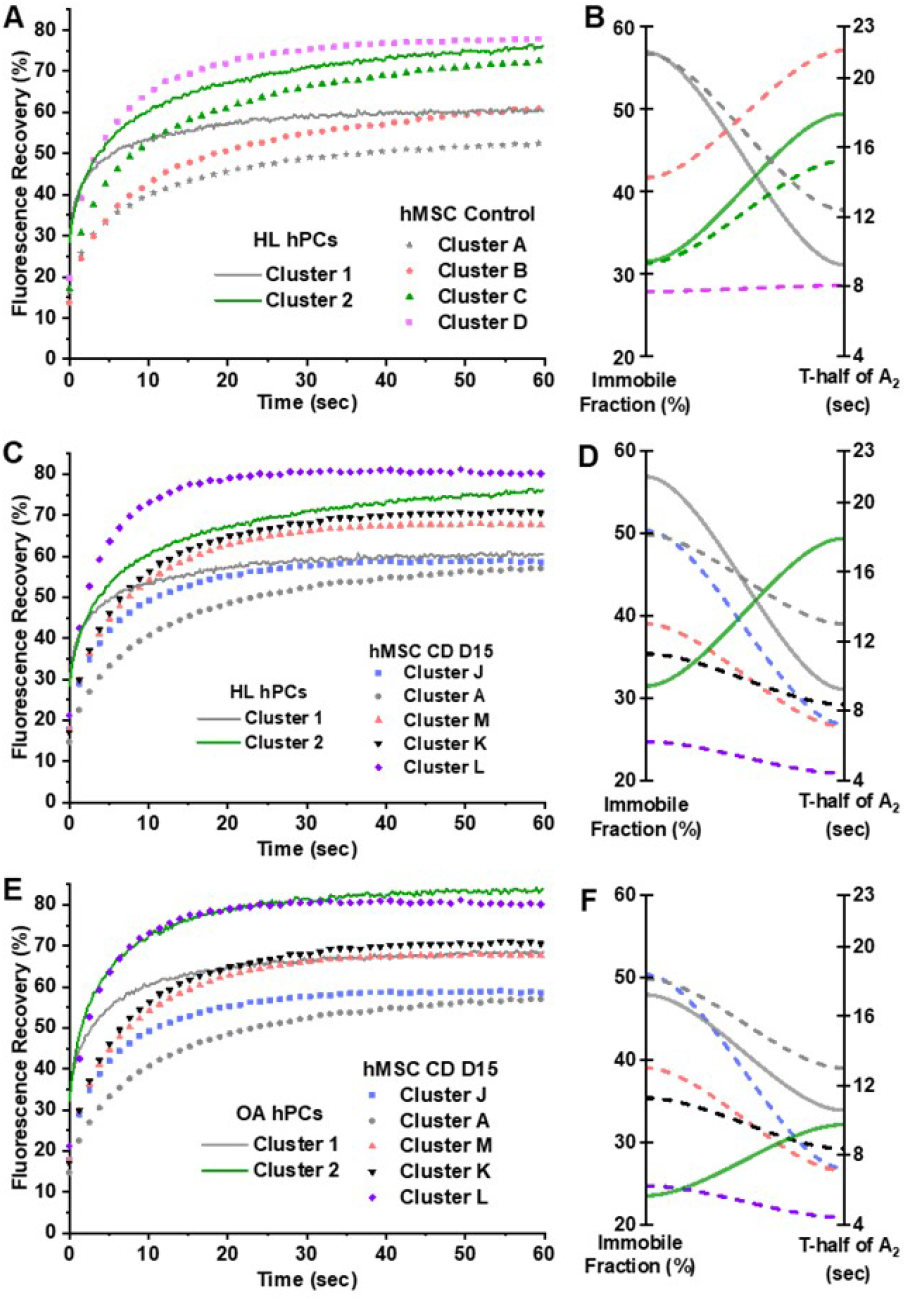
Chondrogenically-differentiated hMSCs may not possess true chondrocyte properties. (A and B) RUNX2 dynamics in undifferentiated hMSCs (control, symbols or dashed lines) comparted to RUNX2 dynamics of healthy (HL) hPCs (continuous line). RUNX2 dynamic rates (IF and t-half of A2) of cluster C of undifferentiated hMSCs were not significantly different from cluster 2 of healthy hPCs. (C and D) RUNX2 dynamics rates of chondrogenically-differentiated hMSCs (day 15, symbols or dashed lines) comparted to RUNX2 dynamics rates of healthy hPCs. RUNX2 dynamic rates (IF and t-half of A1) of cluster J of chondrogenically differentiated hMSCs were not significantly different from cluster 1 of healthy hPCs. (E and F) RUNX2 dynamics rates of chondrogenically-differentiated hMSCs (day 15) comparted to RUNX2 dynamics rates of OA hPCs. RUNX2 dynamic rates of cluster J of chondrogenically-differentiated hMSCs were not significantly different from cluster 1 of healthy hPCs.

Chondrogenic differentiation of hMSCs in 2D culture usually results in hypertrophy (30). TF-FRAP rates of cluster 2 of healthy and OA hPCs did not match with those in any of the clusters of chondrogenically-differentiated hMSCs. Although cluster L of differentiated hMSCs had a close overlap with cluster 2 of OA hPCs, their recovery half-times of A_1_ and A_2_ were significantly different. These data suggest that chondrogenically differentiated hMSCs at day 15 did not form mature hPCs.

### Distinct nuclear localization patterns dictate RUNX2 mobility

We observed differential RUNX2 nuclear localization patterns. After cluster analysis, we found that the cells with lower RUNX2 mobility (i.e. with higher IF and longer recovery half-time) had comparatively higher RUNX2 expression and discrete nuclear localization patterns (Figure 5, cluster A). Cells with higher RUNX2 mobility (i.e. with lower IF and shorter recovery half-times) had comparatively lower RUNX2 expression and diffused nuclear localization patterns (Figure 5, cluster D). Cluster B and C (Figure 5) had similar, but intermediate localization patterns, comparatively less discrete (as compared to cluster A) and less diffused (as compared to cluster D). These distinct localization patterns result in differential RUNX2 dynamics. A montage of nuclei expressing RUNX2 is presented in figure S9-S12 for clusters A-D of undifferentiated hMSCs, respectively. More than 80% of the cells per cluster showed similar nuclear localization patterns.

**Fig. 5.**
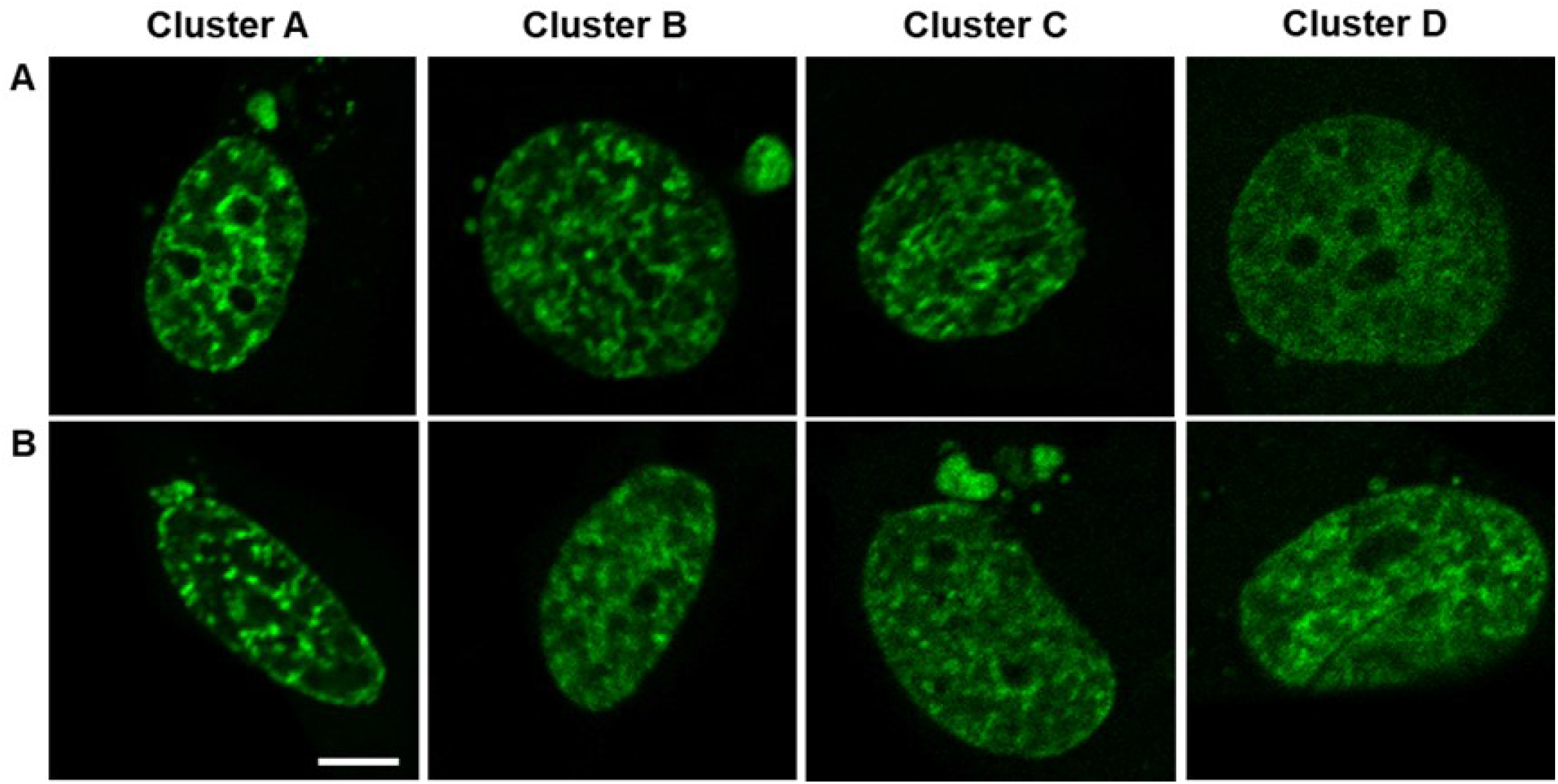
Nuclear localization patterns of eGFP-RUNX2 are distinct among clusters. Nuclei expressing eGFP-RUNX2 from all four clusters of undifferentiated hMSCs are shown here. Most of the cells in each cluster had a nuclear localization pattern as shown in either panel A or B. Cells in cluster A showed higher RUNX2 expression and discrete localization patterns as compared to other clusters, which resulted in lower mobility, i.e., higher immobile fraction and t-half. Cells in cluster D showed lower expression and diffused localization patterns of RUNX2, which resulted in higher mobility, i.e., lower immobile fraction and shorter t-half. Although localization patterns were not different between clusters B and C, cells in these clusters showed intermediate mobility.entiating hMSCs do not respond similarly to the differentiation stimuli. Scale bar: 5*µ*m.

**Fig. 6.**
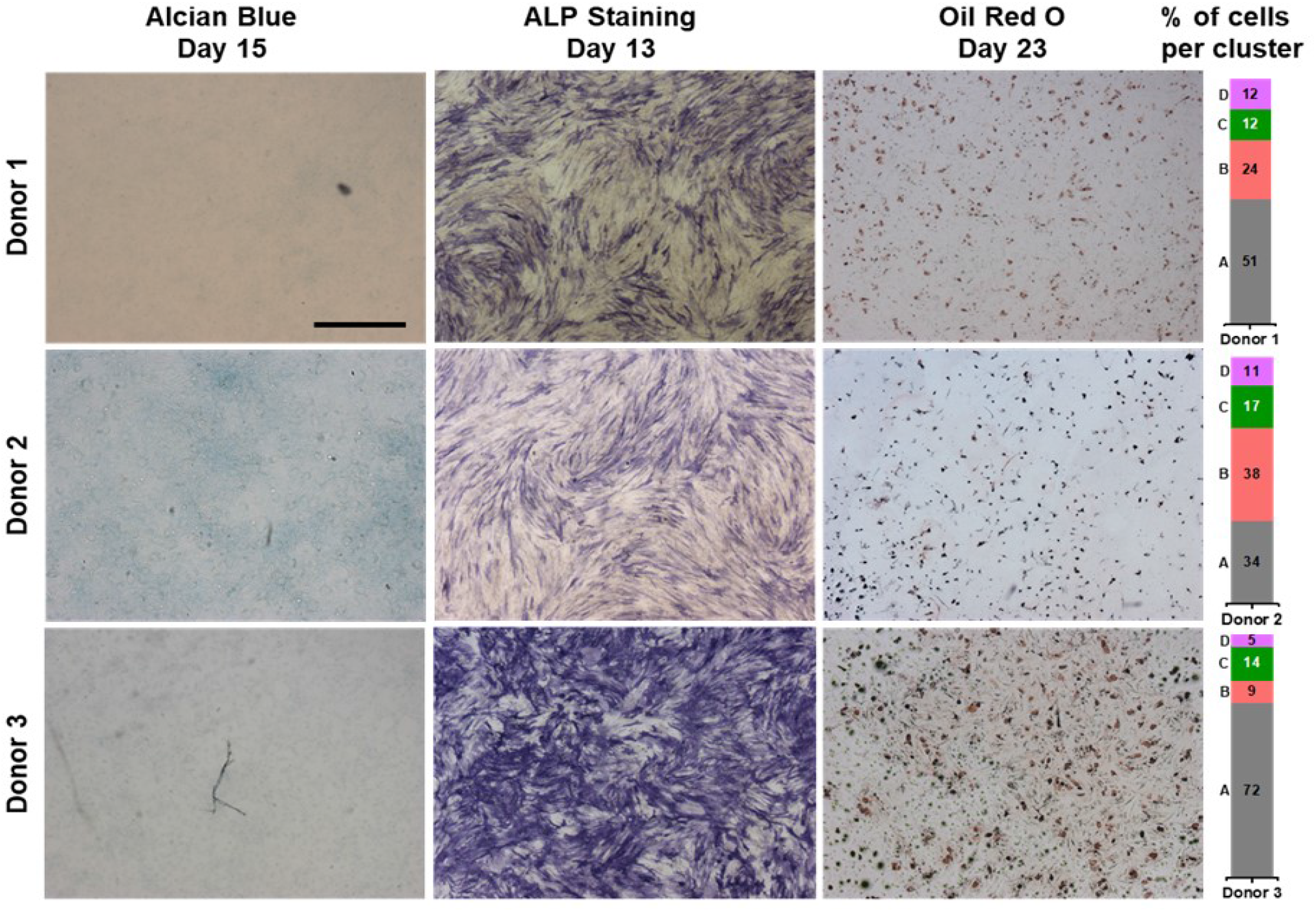
Number of cells present per cluster may determine differentiation potential of a donor. Donor 1 showed poor GAG production and moderate production of ALP and lipid as compared to other donors. In this donor, cluster A contained more than 50% cells and cluster B contained nearly 25% cells. Donor 2 showed high GAG production and moderate ALP and lipid production as compared to other donors. In this donor, nearly 35% of cells were present in clusters A and B. Donor 3 showed poor GAG production, but very high ALP and lipid production as compared to other donors. In this donor, more than 70% of cells were present in the cluster A and clusters B-D together account for remaining 30% cells. Scale bar: 500*µ*m.

Although these differential localization patterns were present in differentiating hMSCs as well, for simplicity, we present here only those observed in undifferentiated cells. To rule out that these localization patterns are due to the difference in cell cycle, we performed cell cycle synchronization study by serum starving the cells for 24 hours. Even after 24 hours of serum starvation and 6 hours after adding serum (post 24-hour starvation), we found at least 4 distinct types of nuclear localization patterns (Figure S13). Although cell cycle synchronization of hMSCs for 24h is shorter as compared to doubling time of hMSCs (30 – 39h), we have observed diverse nuclear localization pattern of RUNX2 in chondrogenically differentiating hMSCs as well, where cell division is limited due to lack of serum in the medium. To check if these localization patterns were the result of over expression of RUNX2, we immuno-stained endogenous RUNX2. Immunofluorescence data also showed that these distinct RUNX2 localization patterns were inherent to hMSC subpopulations (Figure S14).

### Number of cells per subpopulation may determine differentiation potential of a donor

Cluster analysis revealed that the number of cells present per cluster varies per donor. We also observed varied differentiation potential of donors towards a particular lineage. We checked if there is any correlation between the number of cells per cluster and the differentiation potential of a donor. We used three hMSC donors in this study. Interestingly, Donor 1 showed poor chondrogenic potential and comparatively moderate osteogenic and adipogenic potential. From this donor more than 50% cells were found in cluster A and nearly 25% cells in cluster B. Clusters C and D always had <20% cells per cluster in all three donors. Donor 2 showed higher GAG production as compared to other two donors, indicating a higher chondrogenic potential of the donor. This donor showed comparatively moderate ALP and lipid production, indicating moderate differentiation potential towards osteogenic and adipogenic lineage. Of this donor about 35% of cells were found in clusters A and B. Donor 3 showed high osteogenic and adipogenic differentiation potential as evidenced by high ALP and lipid staining at day 23. More than 70% cells of this donor were found in cluster A. These data suggest that the number of progenitor cells per subpopulation of hMSCs may determine the differentiation potential of a donor.

We performed a similar study, in which we determined the mobility of SOX9 in the same donors (16). The cluster IDs of RUNX2 and SOX9 do not correspond. Between SOX9 and RUNX2 studies, we observed a discrepancy in the number of cells present in the different clusters within a donor. This could be due to various reasons. One reason may be that, that including backbone and size of the RUNX2 and SOX9 plasmids were different and they might affect the transfection efficiency of the subpopulations. However, these two independent studies confirm that the number of progenitor cells in subpopulations of hMSCs are not same within a donor, which might determine differentiation potential of a donor.

## Discussion

We have captured changes in the RUNX2 dynamics using TFFRAP at subpopulation level in undifferentiated, chondro-, osteo-, and adipogenically differentiating hMSCs. Cluster analysis of single cell RUNX2 dynamics data indicate that the RUNX2 transcriptional activity may be different among subpopulation of hMSCs. We have shown that number of cells present per subpopulation might determine differentiation potential of a donor towards a particular lineage. Our RUNX2 dynamics study confirms our findings in our previous similar study on SOX9 dynamics in hMSCs (16).

Immobile fraction and recovery half-times of RUNX2 were higher in the initial stages (day 2) of chondrogenic differentiation and gradually reduced at day 15. During osteogenic differentiation, although RUNX2 immobile fraction was lower, many clusters had longer recovery half-times. As expected, recovery half-times were lowest during adipogenic differentiation as compared other two lineages. Despite higher immobile fraction of RUNX2 during adipogenic differentiation, there were no active exchange at the binding sights as indicated by shorter recovery half-times, suggesting no active transcription. These finding are in agreement with previous reports that RUNX2 activity is higher during osteogenic and early stages of chondrogenic differentiation, lower during adipogenic and later stages chondrogenic differentiation (10, 27, 31). The number of subpopulations of cells present in hMSCs are not yet clearly defined (32). Morphology or CD marker based estimations suggest existence of three subpopulations in hMSCs (33). We have previously reported that at least four distinct dynamic rates of SOX9 present in the undifferentiated hMSCs and subsequent FACS analysis showed every subpopulation has distinct SOX9 dynamic rates (16). In line with this report, we have also observed four distinct RUNX2 dynamic rates in undifferentiated hMSCs. We used three hMSC donors for this study and found at least a few cells from every donor in all the four clusters in undifferentiated hMSCs. This further substantiates that hMSCs might have at least four subpopulation of cells, even with the account of donor variation.

Subpopulation of cells respond differently to the same differentiation stimuli. Although all cells were exposed to same environmental stimuli, RUNX2 dynamics did not change in a subpopulation of cells. Interestingly, cluster A in chondrogenic, cluster B in osteogenic and cluster C in adipogenic differentiation were present in all differentiation time points (except cluster C, at day 8 of adipogenic differentiation). This indicates that different subpopulation of cells do not respond to differentiation stimuli of particular lineage and only selected subpopulation of cells can differentiate into a particular lineage. These data are also in line with our previous findings with SOX9 dynamics in hMSCs (16).

Several studies have reported isolation of subpopulation of cells from MSCs based on cell surface CD markers and demonstrated better differentiation of selective subpopulations towards a particular lineage (reviewed, (14)). In our top down approach, we differentiated hMSCs without sorting and performed cluster analysis to identify subpopulations. Interestingly, cluster A of donor 3, which had large number of cells as compared cluster A of other donors, showed comparatively higher osteogenic potential. Our data suggests that number of progenitor cells present per subpopulation of hMSCs is not same among donors and amount of progenitor cells might determine the differentiation potential of a donor towards a particular lineage.

We compared number of clusters present per differentiation lineage and time point between SOX9 (16) and RUNX2. Interestingly, there were same number of clusters in 50% of time points. In the other time points, there were either one cluster higher or lower between SOX9 and RUNX2 (except during adipogenic differentiation at day 23, where the difference was two clusters). Similarity in the number of clusters among differentiation lineages and time points indicate that how these transcription factors function with harmony. Dissimilarity in the number of clusters between SOX9 and RUNX2, per time-point, suggests that changes in dynamics and coupled transcriptional activity of these transcription factors may not change in the same time. In some differentiation time points, it is plausible that two subpopulations might have statistically indistinguishable dynamic rates and they were considered as single cluster by the clustering algorithm. This could explain the difference in the number of subpopulations (or clusters) between SOX9 and RUNX2, within a time-point.

Presence of lipid droplets allowed us to identify adipogenically differentiated hMSCs (at day 23, without Oil Red O staining) during TF-FRAP measurements. We expected all adipogenically-differentiated cells would have same RUNX2 dynamics. Surprisingly, these cells were also distributed across all clusters with differential dynamics rates (except in the cluster I). ALP and Oil Red O staining also show varied ALP and lipid droplet production at the single cell level in differentiated cells. This suggests that although a particular subpopulation of hMSCs may have higher differentiation potential towards a particular lineage differentiation, cells from other subpopulations might also respond and differentiate into other lineages. However, their molecular properties might be different and may not bear the true molecular signature of the differentiated cell type. This observation is in line with the findings by Leyva-Leyva et al (2012), they isolated CD105+ and CD105– subpopulations from MSCs and demonstrated that although both the subpopulations can osteogenically differentiate, the former had higher differentiation potential as compared to the later (34).

RUNX2 activity is essential for chondrocyte differentiation and survival (35). TF-FRAP identified cluster C (with lower IF, but comparatively longer recovery half-times) as a subpopulation with chondrogenic potential. However, 2D chondrogenic differentiation hMSCs did not possess the properties of true articular chondrocytes. Thus, RUNX2 dynamic rates of at least one of the clusters in healthy and OA hPCs did not match with clusters in chondrogenically-differentiated hMSCs. This observation is in line with our SOX9 dynamics study that chondrogenically-differentiated hMSCs did not form mature chondrocytes (16).

We have previously reported that the SOX9 dynamics is coupled to its nuclear localization patterns (16). As expected, we observed that the RUNX2 also show differential nuclear localization patterns among subpopulation of cells and which dictates its dynamics. Higher expression and discrete localization patterns always resulted in a lower mobility and vice versa. It is interesting to note that differential localization patterns and coupled dynamics are consistent among transcription factors. Previous studies have shown that presence of discrete sub-nuclear foci of transcription factors indicate increased transcriptional output (36–38) and Zaidi et al (2001) has shown this specifically for RUNX2 (39). Together with these reports, the cluster A of undifferentiated hMSCs, which shows discrete sub-nuclear foci coupled lower mobility, might be the subpopulation with higher osteogenic potential.

It is already known that there is a significant difference in the RUNX2 target gene expression profile among subpopulations of hMSCs (40). Subpopulations with more chondrogenic and osteogenic differentiation potential expressing higher SOX9 and RUNX2 target genes respectively, have been described already (34, 41, 42). This suggests that SOX9 and RUNX2 transcriptional activity is different among hMSC subpopulations. We have previously shown that changes in SOX9 dynamics as an indicator of changes in its transcriptional activity (19). Together, these findings indicate that RUNX2 transcriptional activity might be different among these subpopulations due to their differential dynamics, expression and nuclear localization pattern.

We have demonstrated how RUNX2 transcriptional activity change at the subpopulation level during multi-lineage differentiation of hMSCs. Moreover, our findings on RUNX2 and SOX9 (16) dynamics indicate that the transcription factor dynamics is coupled to their nuclear localization pattern. In addition, TF-FRAP method can be used to define the number of subpopulations in heterogenic cells and to determine the differentiation potential of a donor towards a particular lineage.

