## Supplementary material for "Mapping RUNX2 transcriptional dynamics during multi-lineage differentiation of human mesenchymal stem cells": Figure S1

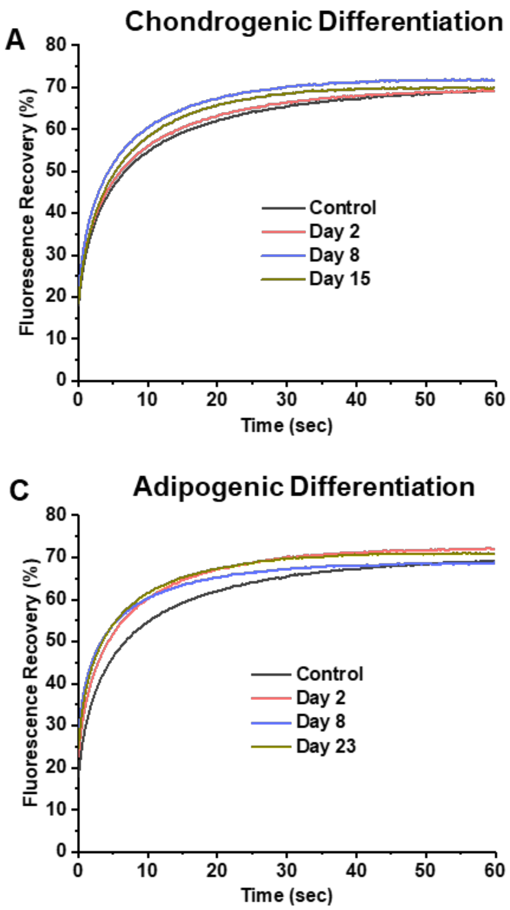


***Figure S1. A-C. Averaged FRAP recovery curves per time point shows the RUNX2 mobility only increased during any differentiation lineage as compared to the undifferentiated control.***
