## Supplementary material for "Mapping RUNX2 transcriptional dynamics during multi-lineage differentiation of human mesenchymal stem cells": Figure S2

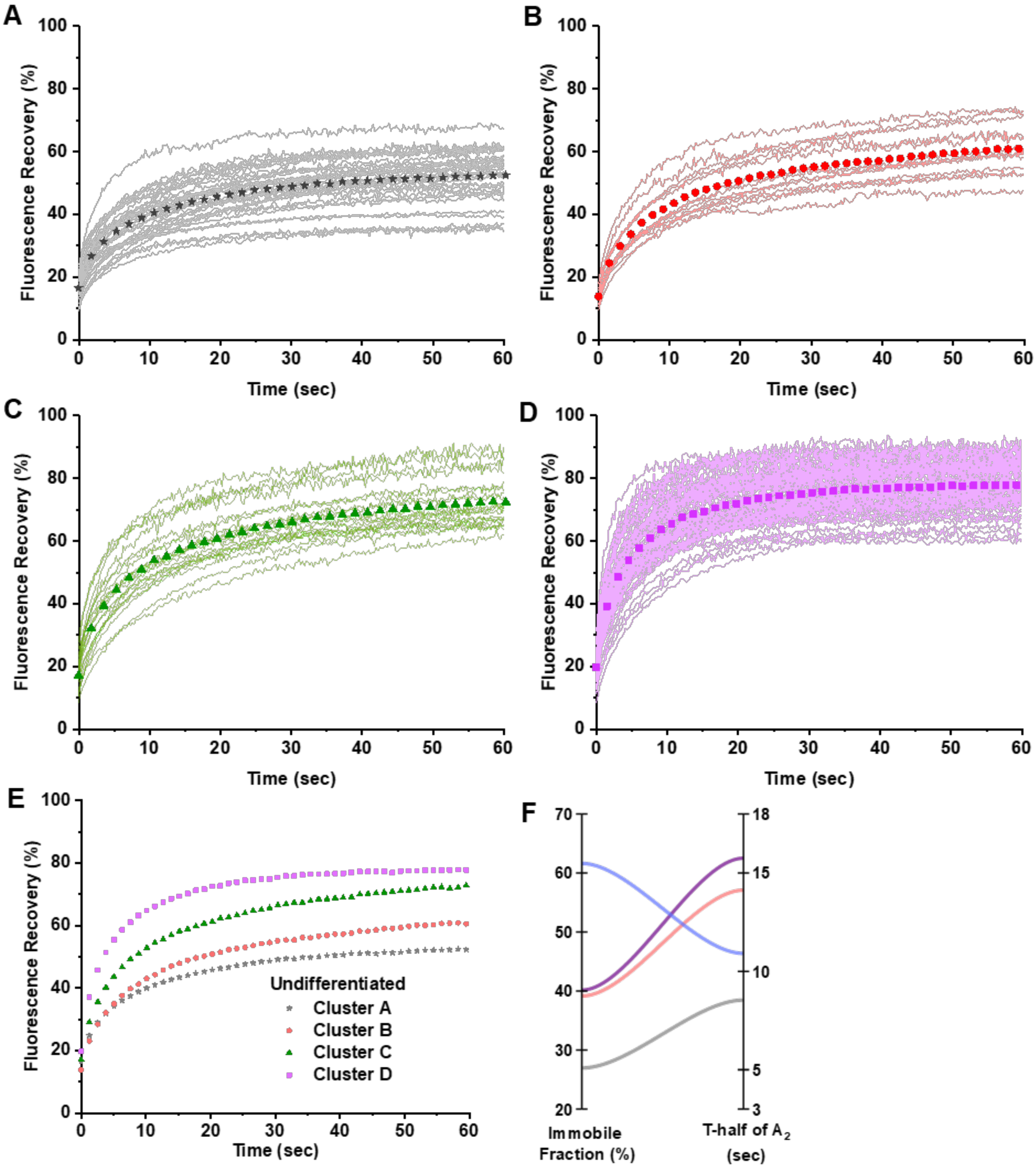


***Figure S2. FRAP curves segregated by unsupervised hierarchical clustering show at least four types of RUNX2 mobility pattern in the heterogenic population of hMSCs.*** *(A-D) RUNX2 mobility pattern in the individual cells of cluster 1 to 4 respectively. E. Average of FRAP curves per cluster. F. Parallel plot show the changes and the relationship between immobile fraction and recovery half-time of $A_2$ in these clusters (n = 137).*
