## Supplementary material for "Mapping RUNX2 transcriptional dynamics during multi-lineage differentiation of human mesenchymal stem cells": Figure S3

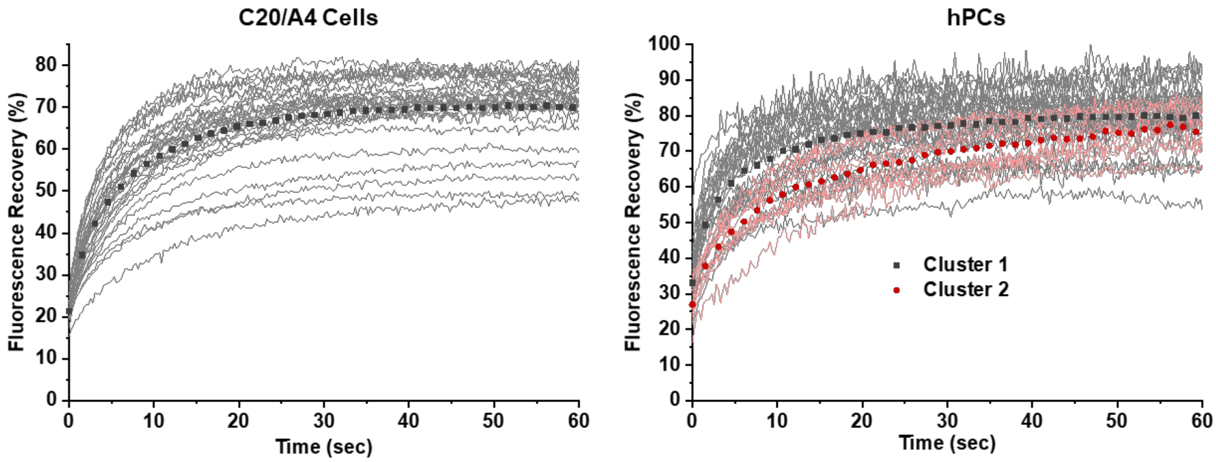


***Figure S3. FRAP cures of eGFP-RUNX2 in C20/A4 cells (A) show only one type of mobility pattern and human primary chondrocytes isolated from OA joint (B) show two clusters of mobility pattern.***
