## Supplementary material for "Mapping RUNX2 transcriptional dynamics during multi-lineage differentiation of human mesenchymal stem cells": Figure S4

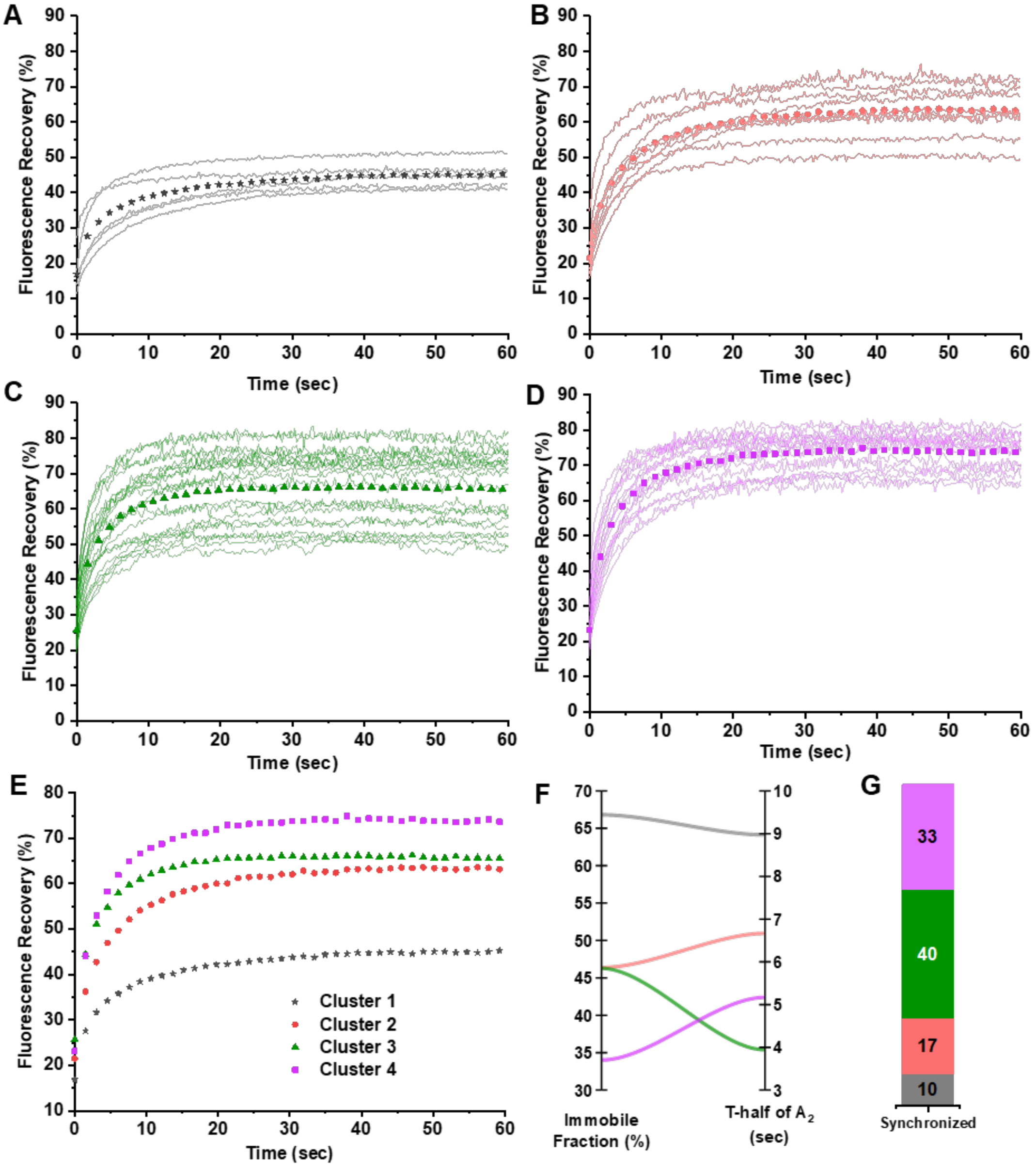


***Figure S4. eGFP-RUNX2 dynamics in the control group of hMSCs after synchronization for 24 hours show that the differential SOX9 dynamics is not due to the cell cycle.*** *If the differential RUNX2 mobility patters were due to cell cycle, synchronization study will show less than four clusters. (A) FRAP curves of RUNX2 show the mobility pattern among the clusters, (B) parallel plot showing the relationship between immobile fraction and t-half of A2 among the clusters and (C) the stacked bar chart showing the percentage of cells per cluster (n = 60).*
