## Supplementary material for "Mapping RUNX2 transcriptional dynamics during multi-lineage differentiation of human mesenchymal stem cells": Figure S6

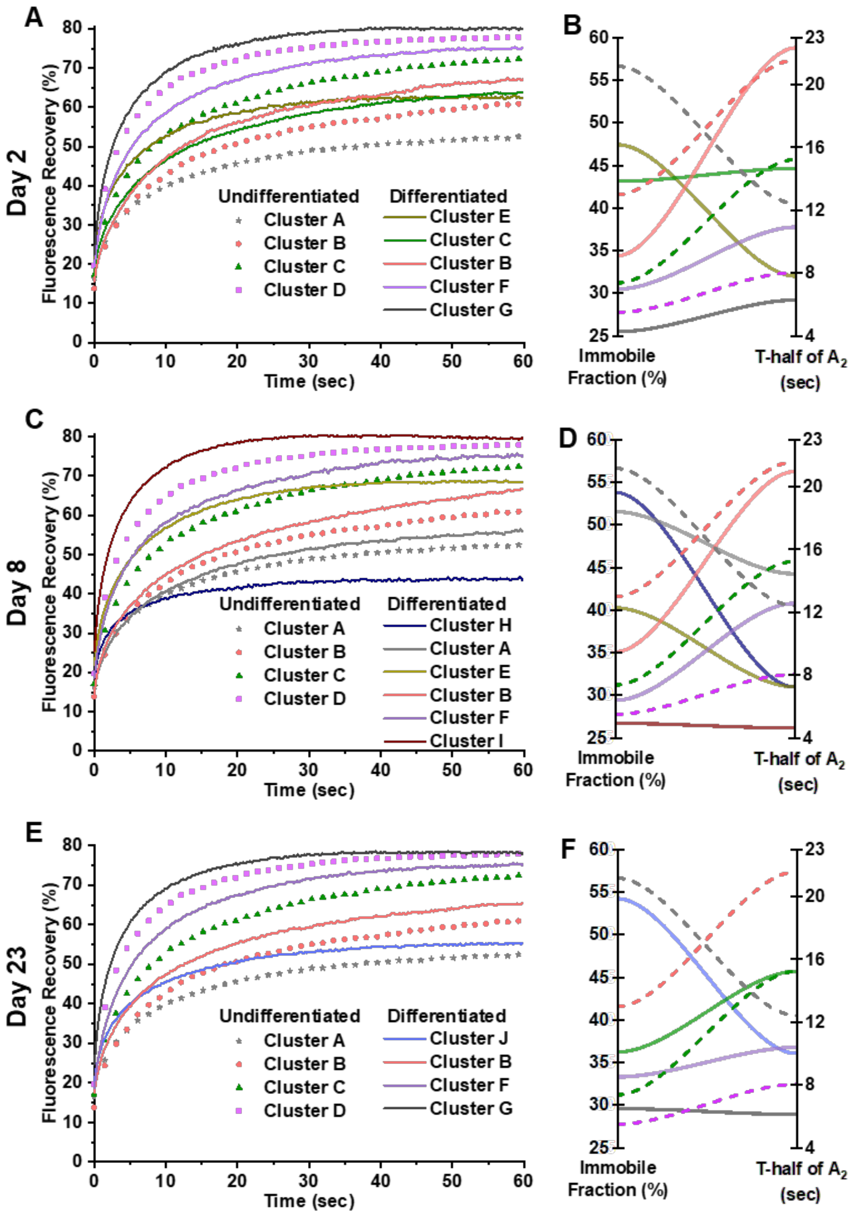


***Figure S6. Mobility of eGFP-RUNX2 in undifferentiated and osteogenically differentiating hMSCs as measured by FRAP.*** *In the differentiated group (continuous line), five clusters with distinct dynamic rates appeared at all three time points. FRAP recovery curves on the left (A, C and E) show changes in the mobility pattern of eGFP-RUNX2 in the osteogenically differentiating clusters as compared to clusters in undifferentiated cells (symbols or dash line) for day 2, 8 and 23, respectively. Parallel plot on the right (B, D and F) show the relationship between IF and t½ of* $A_2$ *and its changes between undifferentiated (dashed line) and differentiating hMSCs (continuous line) for day 2, 8 and 23, respectively. Day 2, 8 and 23 had five, six and four clusters, respectively. Three hMSC donors were used for this study. Combined data from all the donors are shown here. N ≥ 119, per time point.*
