## Supplementary material for "Mapping RUNX2 transcriptional dynamics during multi-lineage differentiation of human mesenchymal stem cells": Figure S7

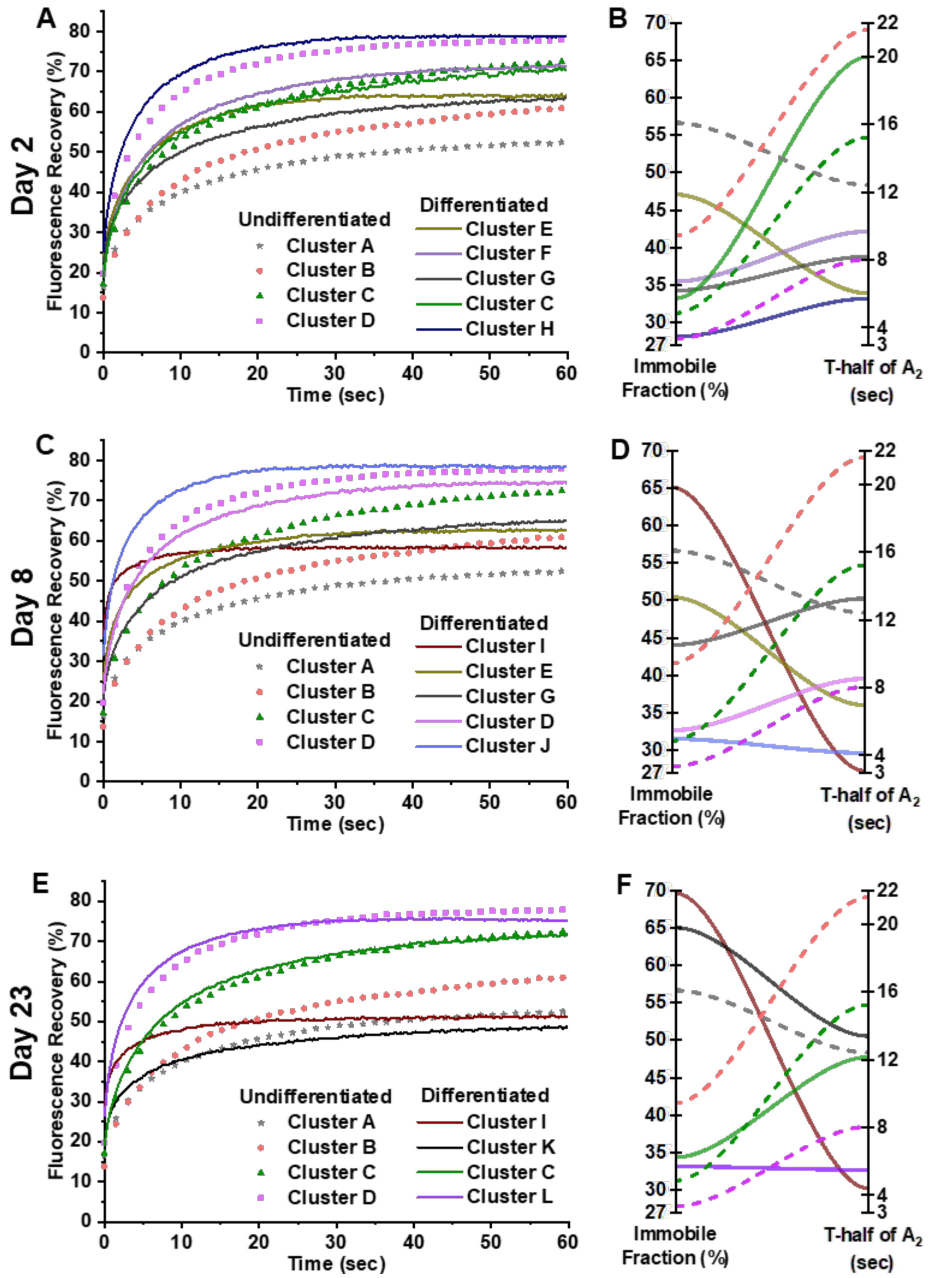


***Figure S7. Mobility of eGFP-RUNX2 in undifferentiated and adipogenically differentiating hMSCs as measured by FRAP.*** *In the differentiated group (continuous line), three clusters with distinct dynamic rates appeared at day 2 (A and B), day 8 (C and D) and day 23 (E and F). FRAP recovery curves on the left show changes in the mobility pattern of eGFP-RUNX2 in the adipogenically differentiating clusters as compared to clusters in undifferentiated cells (symbols). Parallel plot on the right show the relationship between IF and t½ of $A_2$ and its changes between undifferentiated (continuous line) and differentiating hMSCs (dashed line). Day 2 and 8 had five clusters each and day 23 had four clusters. Three hMSC donors were used for this study. Combined data from all the donors are shown here. N ≥ 126, per time point.*
