## Supplementary material for "Mapping RUNX2 transcriptional dynamics during multi-lineage differentiation of human mesenchymal stem cells": Figure S8

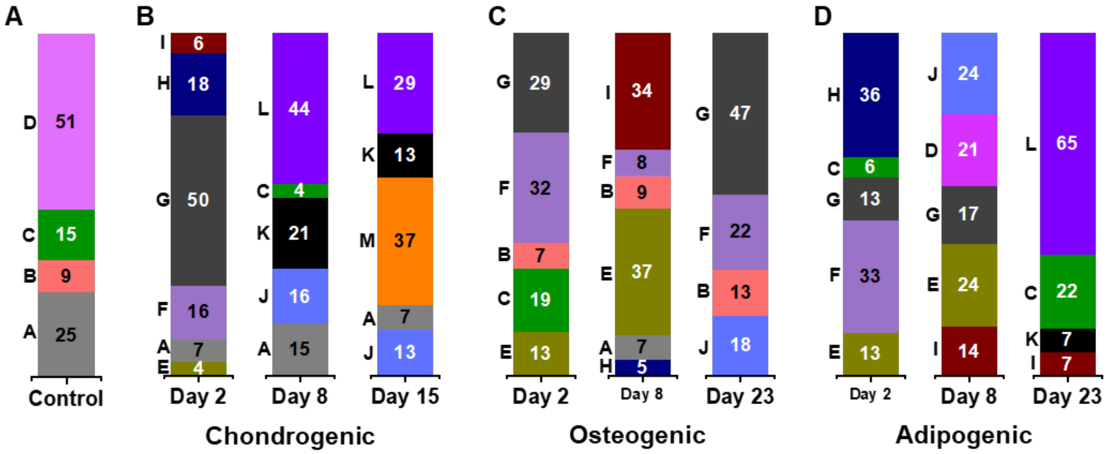


***Figure S8. Percentage of cells present in each cluster is variable depending on the differentiation lineage and time.*** *Bar graphs show proportion of cells in present in each cluster of the undifferentiated hMSCs (A), chondrogenically (B), osteogenically (C) and adipogenically (D) differentiating hMSCs. Letters to the left of the bar graph indicate the cluster ID.*
