## Supplementary material for "Mapping RUNX2 transcriptional dynamics during multi-lineage differentiation of human mesenchymal stem cells": Figure S9

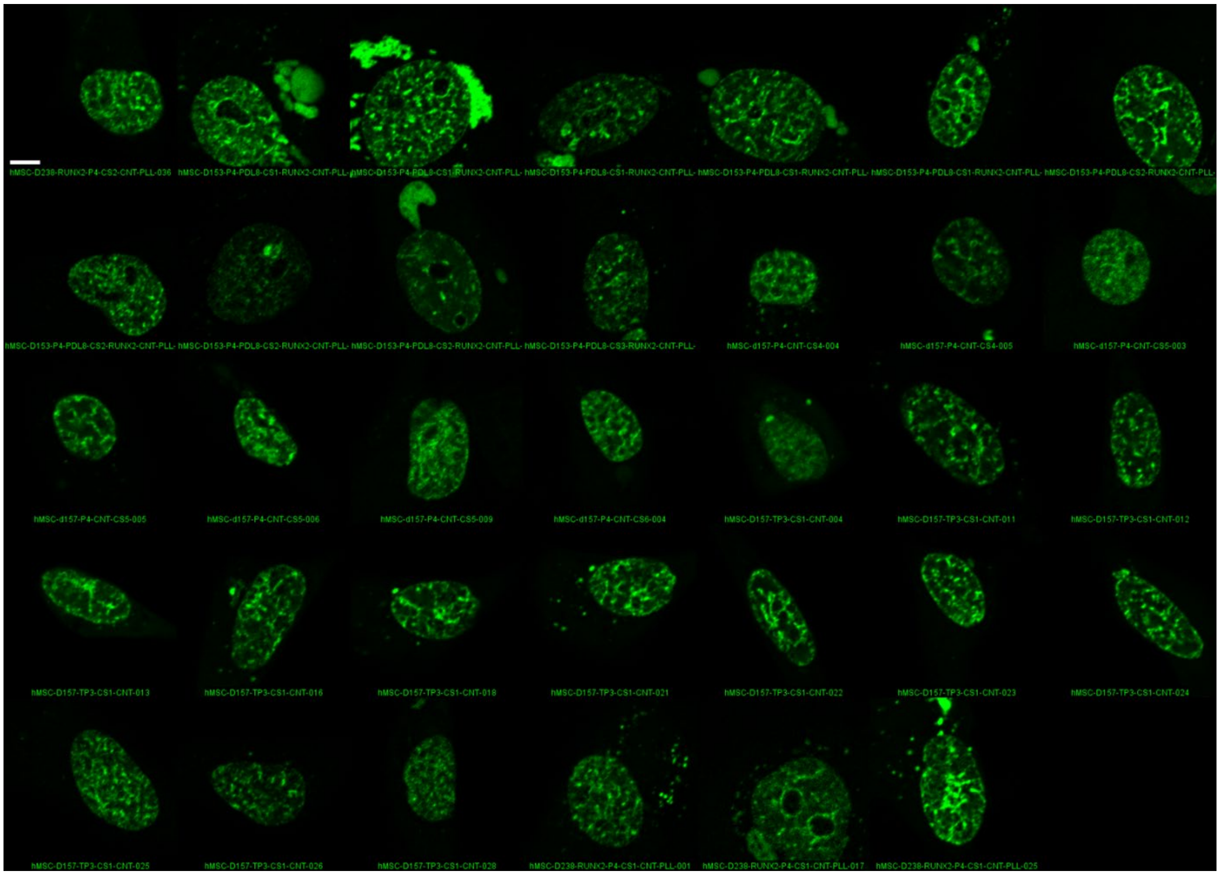


***Figure S9. Montage of nuclei showing nuclear localization pattern of eGFP-RUNX2 in the cluster A of control group (undifferentiated hMSCs).*** *Scale bar: 5 µm.*
