## Supplementary material for "Mapping RUNX2 transcriptional dynamics during multi-lineage differentiation of human mesenchymal stem cells": Figure S12

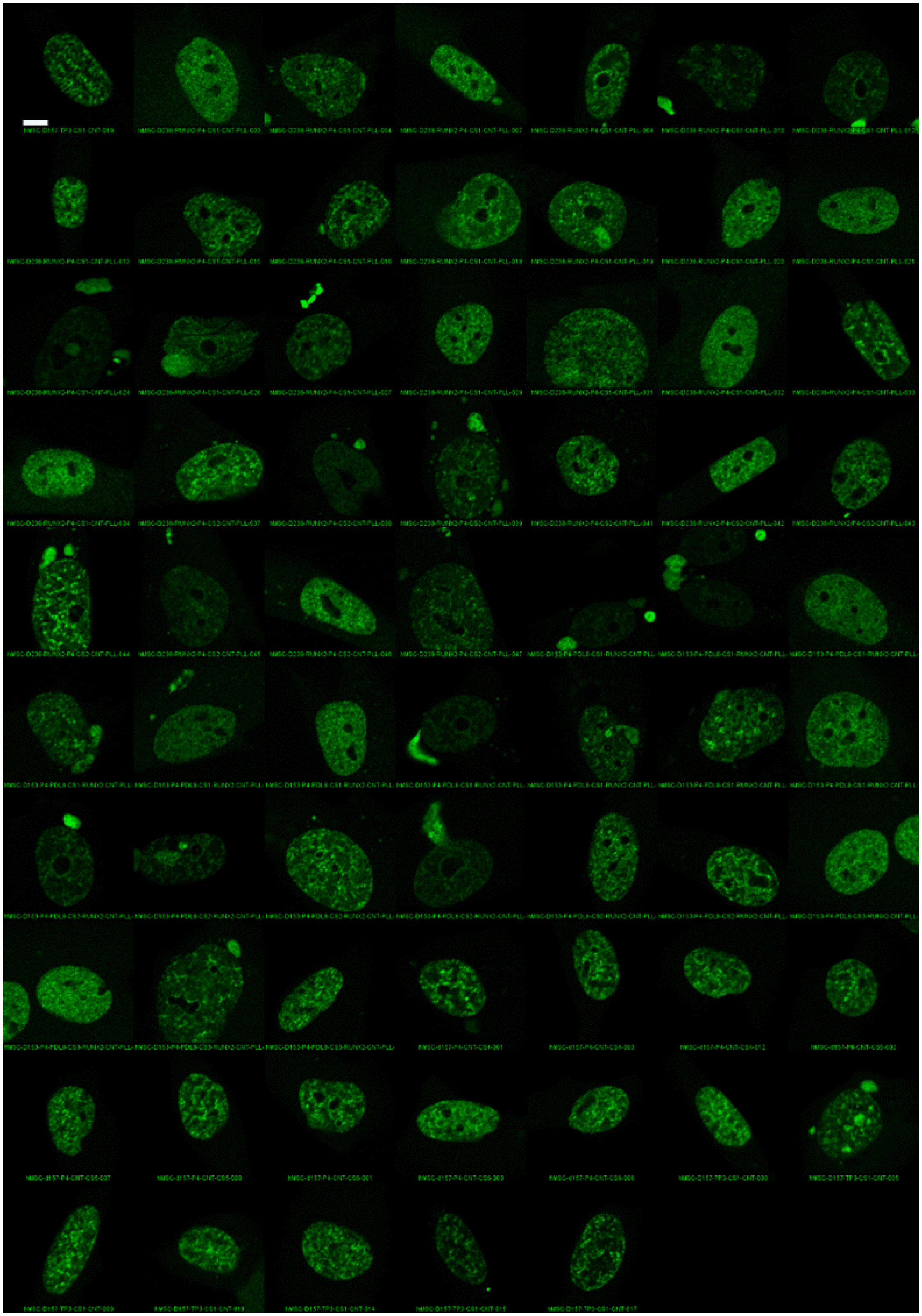

***Figure S12. Montage of nuclei showing nuclear localization pattern of eGFP-RUNX2 in the cluster D of control group (undifferentiated hMSCs).*** *Scale bar: 5 µm.*
