## Supplementary material for "Mapping RUNX2 transcriptional dynamics during multi-lineage differentiation of human mesenchymal stem cells": Figure S14

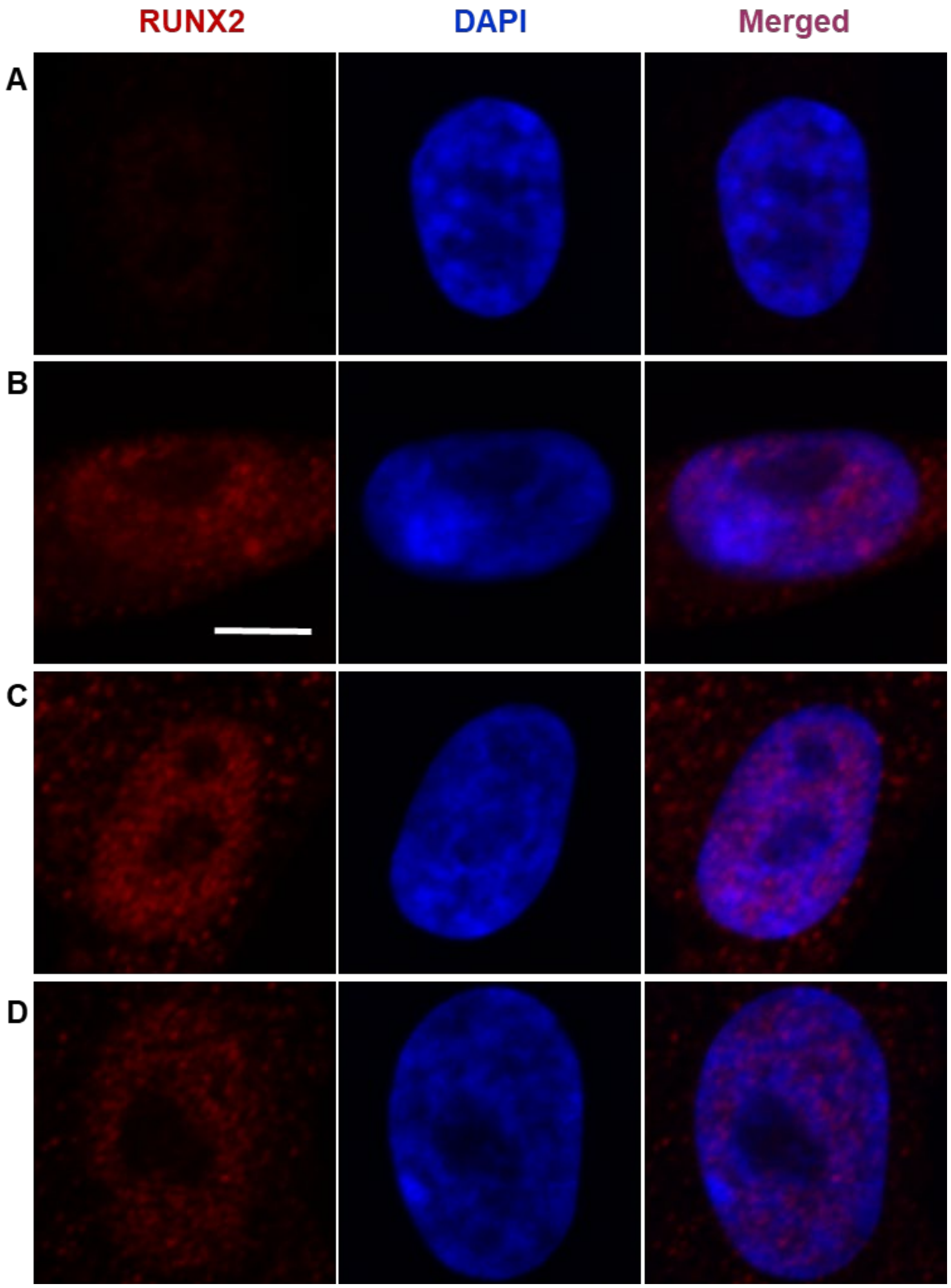


***Figure S14. Immunostaining of endogenous RUNX2 in hMSCs show at least four types nuclear localization patterns in the undifferentiated hMSCs.*** *This indicates that the differential nuclear localization patterns of RUNX2 in the eGFP-RUNX2 transfected cells are not due to its overexpression. Scale bar: 5 µm.*
