## Supplementary material for "Mapping RUNX2 transcriptional dynamics during multi-lineage differentiation of human mesenchymal stem cells": Figure S15

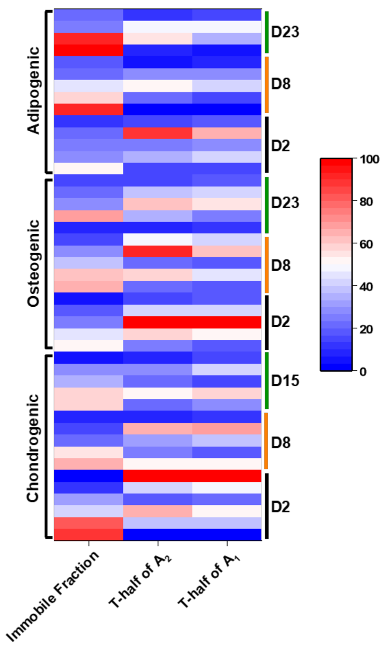


***Figure S15. Heat map comparing FRAP rates of eGFP-RUNX2 during chondro-, osteo-, and adipogenic differentiation, averaged at subpopulation level.*** *Clusters in osteogenic differentiation show longer Recovery half-times as compared other differentiation lineages. Clusters in chondrogenic differentiation show shorter recovery half-times as compared to osteogenic differentiation. Clusters in adipogenic differentiation show shorter recovery half-times as compared to other two differentiation lineages. Immobile fraction was higher in the initial stages chondrogenic differentiation and later stages on adipogenic differentiation.*
