## Supplementary material for "Mapping RUNX2 transcriptional dynamics during multi-lineage differentiation of human mesenchymal stem cells": Table S1

***Table S1. FRAP rates of eGFP-RUNX2, number and percentage of cells in the individual clusters of undifferentiated hMSCs.***

| | **Immobile Fraction (%)** | **T-half of $A_2$ (sec)** | **T-half of $A_1$ (sec)** | **No. of cells** | **Percentage of cells** |
|---|---|---|---|---|---|
| Cluster A | 56.6 ± 9.0 | 12.38 ± 3.38 | 1.53 ± 0.38 | 34 | 25 |
| Cluster B | 41.5 ± 10.0 | 21.55 ± 3.99 | 3.14 ± 0.54 | 13 | 9 |
| Cluster C | 31.1 ± 9.3 | 15.20 ± 2.16 | 2.10 ± 0.45 | 20 | 15 |
| Cluster D | 27.7 ± 9.0 | 8.00 ± 2.52 | 1.34 ± 0.48 | 70 | 51 |
