## Supplementary material for "Mapping RUNX2 transcriptional dynamics during multi-lineage differentiation of human mesenchymal stem cells": Table S2

***Table S2. FRAP rates of eGFP-RUNX2, number and percentage of cells in the individual clusters of chondrogenically differentiating hMSCs. If same cluster is found more than two times, * indicates the significance pair.***

| | | Immobile Fraction (%) | T-half of $A_2$ (sec) | T-half of $A_1$ (sec) | No. of cells | % of cells | P-Value | | |
|---|---|---|---|---|---|---|---|---|---|
| | | | | | | | IF | $A_2$ | $A_1$ |
| **Day 2** | Cluster E | 63.9 ± 7.0 | 3.40 ± 0.15 | 0.40 ± 0.09 | 5 | 4 | | | |
| | Cluster A | 61.2 ± 7.4 | 10.24 ± 3.22 | 1.60 ± 0.26 | 8 | 7 | 0.11 | 0.09 | 0.84 |
| | Cluster F | 41.9 ± 6.3 | 15.50 ± 3.91 | 2.04 ± 0.51 | 19 | 16 | | | |
| | Cluster G | 36.6 ± 8.7 | 6.37 ± 1.79 | 1.04 ± 0.32 | 60 | 50 | | | |
| | Cluster H | 27.5 ± 3.7 | 11.16 ± 2.42 | 1.93 ± 0.43 | 22 | 18 | | | |
| | Cluster I | 22.3 ± 7.1 | 22.37 ± 4.85 | 3.62 ± 0.38 | 7 | 6 | | | |
| **Day 8** | Cluster A* | 53.1 ± 6.7 | 12.73 ± 1.88 | 2.00 ± 0.40 | 19 | 15 | 0.23 | 0.66 | 0.001 |
| | Cluster J | 47.9 ± 10.4 | 7.63 ± 1.78 | 0.94 ± 0.37 | 20 | 16 | | | |
| | Cluster K | 32.4 ± 7.0 | 8.97 ± 1.55 | 1.60 ± 0.29 | 26 | 21 | | | |
| | Cluster C | 28.7 ± 9.0 | 15.88 ± 2.76 | 2.61 ± 0.34 | 5 | 4 | 0.40 | 0.52 | 0.03 |
| | Cluster L | 25.7 ± 4.4 | 4.54 ± 1.19 | 0.80 ± 0.22 | 55 | 44 | | | |
| **Day 15** | Cluster J | 50.2 ± 5.7 | 7.20 ± 1.30 | 1.26 ± 0.44 | 17 | 13 | 0.14 | 0.40 | 0.02 |
| | Cluster A* | 49.7 ± 4.4 | 12.99 ± 1.75 | 2.27 ± 0.40 | 9 | 7 | 0.18 | 0.69 | 0.09 |
| | Cluster M | 38.9 ± 5.1 | 7.07 ± 1.59 | 0.91 ± 0.25 | 47 | 37 | | | |
| | Cluster K | 35.3 ± 4.9 | 8.32 ± 0.92 | 1.76 ± 0.33 | 16 | 13 | 0.06 | 0.31 | 0.16 |
| | Cluster L | 24.6 ± 3.9 | 4.37 ± 1.11 | 0.83 ± 0.27 | 37 | 29 | 0.20 | 0.41 | 0.97 |
