## Supplementary material for "Mapping RUNX2 transcriptional dynamics during multi-lineage differentiation of human mesenchymal stem cells": Table S3

***Table S3. FRAP rates of SOX9-mGFP, number and percentage of cells in the individual clusters of osteogenically differentiating hMSCs. If same cluster is found more than two times, * and # indicate the significance pair.***

| | | Immobile Fraction (%) | T-half of $A_2$ (sec) | T-half of $A_1$ (sec) | No. of cells | % of cells | P-Value | | |
|---|---|---|---|---|---|---|---|---|---|
| | | | | | | | IF | $A_2$ | $A_1$ |
| **Day 2** | Cluster E | 47.4 ± 6.6 | 7.76 ± 2.41 | 0.97 ± 0.38 | 16 | 13 | | | |
| | Cluster C | 43.1 ± 5.5 | 14.63 ± 2.77 | 2.09 ± 0.32 | 23 | 19 | 0.00001 | 0.43 | 0.89 |
| | Cluster B | 34.3 ± 11.6 | 22.31 ± 4.73 | 3.53 ± 0.51 | 9 | 7 | 0.16 | 0.64 | 0.18 |
| | Cluster F$^{\#}$* | 30.4 ± 6.1 | 10.89 ± 2.33 | 1.67 ± 0.38 | 40 | 32 | | | |
| | Cluster G | 25.4 ± 5.4 | 6.24 ± 1.96 | 0.97 ± 0.31 | 36 | 29 | | | |
| **Day 8** | Cluster H | 53.8 ± 8.1 | 7.21 ± 2.80 | 1.01 ± 0.33 | 6 | 5 | | | |
| | Cluster A | 51.5 ± 5.2 | 14.42 ± 3.19 | 1.84 ± 0.44 | 9 | 7 | 0.18 | 0.14 | 0.08 |
| | Cluster E | 40.2 ± 5.9 | 7.22 ± 2.06 | 0.99 ± 0.34 | 47 | 37 | 0.00001 | 0.40 | 0.86 |
| | Cluster B* | 35.0 ± 8.1 | 20.92 ± 3.38 | 2.35 ± 0.30 | 12 | 9 | 0.07 | 0.89 | 0.00001 |
| | Cluster F$^{\#}$ | 29.3 ± 6.0 | 12.52 ± 2.77 | 1.77 ± 0.31 | 10 | 8 | 0.88 | 0.12 | 0.30 |
| | Cluster I | 26.6 ± 6.0 | 4.59 ± 1.68 | 0.72 ± 0.34 | 43 | 34 | | | |
| **Day 23** | Cluster J | 54.2 ± 11.9 | 9.97 ± 2.87 | 1.22 ± 0.34 | 21 | 18 | | | |
| | Cluster B* | 36.2 ± 7.9 | 15.20 ± 6.49 | 2.18 ± 0.76 | 16 | 13 | 0.73 | 0.003 | 0.32 |
| | Cluster F* | 33.2 ± 10.1 | 10.35 ± 3.08 | 1.68 ± 0.26 | 26 | 22 | 0.30 | 0.13 | 0.68 |
| | Cluster G | 29.5 ± 8.6 | 6.08 ± 2.34 | 0.94 ± 0.32 | 56 | 47 | 0.02 | 0.32 | 0.37 |
