## Supplementary material for "Mapping RUNX2 transcriptional dynamics during multi-lineage differentiation of human mesenchymal stem cells": Table S4

***Table S4. FRAP rates of eGFP-RUNX2, number and percentage of cells in the individual clusters of adipogenically differentiating hMSCs. If same cluster is found more than two times, * indicate the significance pair.***

| | | Immobile Fraction (%) | T-half of $A_2$ (sec) | T-half of $A_1$ (sec) | No. of cells | % of cells | P-Value | | |
|---|---|---|---|---|---|---|---|---|---|
| | | | | | | | IF | $A_2$ | $A_1$ |
| **Day 2** | Cluster E | 47.0 ± 5.0 | 6.01 ± 1.39 | 0.82 ± 0.35 | 15 | 13 | | | |
| | Cluster F | 35.4 ± 6.4 | 9.64 ± 2.47 | 1.71 ± 0.42 | 39 | 33 | | | |
| | Cluster G | 34.2 ± 12.3 | 8.15 ± 3.48 | 1.27 ± 0.52 | 15 | 13 | | | |
| | Cluster C* | 33.1 ± 9.7 | 19.95 ± 4.17 | 2.44 ± 0.66 | 7 | 6 | 0.60 | 0.01 | 0.21 |
| | Cluster H | 28.0 ± 6.4 | 5.68 ± 1.64 | 0.96 ± 0.34 | 43 | 36 | | | |
| **Day 8** | Cluster I | 65.0 ± 13.0 | 3.05 ± 0.97 | 0.37 ± 0.09 | 17 | 14 | | | |
| | Cluster E | 50.3 ± 9.8 | 6.94 ± 1.73 | 0.87 ± 0.14 | 29 | 24 | 0.39 | 0.12 | 0.40 |
| | Cluster G | 43.9 ± 14.7 | 13.23 ± 2.24 | 1.78 ± 0.29 | 20 | 17 | 0.56 | 0.18 | 0.83 |
| | Cluster D | 32.5 ± 10.5 | 8.50 ± 1.56 | 1.29 ± 0.29 | 25 | 21 | 0.04 | 0.47 | 0.92 |
| | Cluster J | 31.4 ± 5.4 | 4.11 ± 1.02 | 0.65 ± 0.20 | 28 | 24 | | | |
| **Day 23** | Cluster I | 69.5 ± 8.9 | 4.38 ± 1.54 | 0.56 ± 0.23 | 9 | 7 | 0.33 | 0.07 | 0.01 |
| | Cluster K | 64.9 ± 5.2 | 13.37 ± 3.87 | 1.51 ± 0.40 | 9 | 7 | | | |
| | Cluster C* | 34.3 ± 11.0 | 12.11 ± 4.20 | 1.97 ± 0.38 | 28 | 22 | 0.79 | 0.00001 | 0.11 |
| | Cluster L | 33.0 ± 9.1 | 5.46 ± 1.82 | 0.82 ± 0.28 | 84 | 65 | | | |
