## Supplementary material for "Mapping RUNX2 transcriptional dynamics during multi-lineage differentiation of human mesenchymal stem cells": Table S5

***Table S5. eGFP-RUNX2 FRAP rates of undifferentiated hMSCs and chondrogenically differentiated hMSCs (at day 15) are compared to healthy and OA hPCs. Symbols in superscript indicate significance pair.***

| | | Immobile Fraction (%) | T-half of $A_2$ (sec) | T-half of $A_1$ (sec) | P-Value | | |
|---|---|---|---|---|---|---|---|
| | | | | | IF | $A_2$ | $A_1$ |
| **hMSC Control** | Cluster A | 56.6 ± 9.0 | 12.38 ± 3.38 | 1.53 ± 0.38 | | | |
| | Cluster B | 41.5 ± 10.0 | 21.55 ± 3.99 | 3.14 ± 0.54 | | | |
| | Cluster C* | 31.1 ± 9.3 | 15.20 ± 2.16 | 2.10 ± 0.45 | | | |
| | Cluster D | 27.7 ± 9.0 | 8.00 ± 2.52 | 1.34 ± 0.48 | | | |
| **hMSC CD D15** | Cluster J$^{\$\#}$ | 50.2 ± 5.7 | 7.20 ± 1.30 | 1.26 ± 0.44 | | | |
| | Cluster A | 49.7 ± 4.4 | 12.99 ± 1.75 | 2.27 ± 0.40 | | | |
| | Cluster M | 38.9 ± 5.1 | 7.07 ± 1.59 | 0.91 ± 0.25 | | | |
| | Cluster K | 35.3 ± 4.9 | 8.32 ± 0.92 | 1.76 ± 0.33 | | | |
| | Cluster L | 24.6 ± 3.9 | 4.37 ± 1.11 | 0.83 ± 0.27 | | | |
| **hPCs** | HL Cluster 1$^{\$}$ | 56.8 ± 12.4 | 9.22 ± 2.07 | 0.97 ± 0.30 | 0.11 | 0.006 | 0.054 |
| | HL Cluster 2* | 31.3 ± 11.8 | 17.90 ± 7.90 | 1.83 ± 0.51 | 0.60 | 0.60 | 0.049 |
| | OA Cluster 1$^{\#}$ | 47.8 ± 8.2 | 10.55 ± 4.97 | 1.44 ± 0.59 | 0.45 | 0.13 | 0.40 |
| | OA Cluster 2 | 23.4 ± 6.5 | 9.70 ± 5.95 | 1.23 ± 0.50 | | | |
